# Proximity labelling and evolutionary evidence reveal insights into IL-1α nuclear networks and function

**DOI:** 10.1101/2023.06.26.546506

**Authors:** Rose Wellens, Victor S. Tapia, Hayley Bennett, Antony Adamson, Jack Rivers-Auty, Jack P. Green, Gloria Lopez-Castejon, David Brough, Christopher Hoyle

## Abstract

Interleukin (IL)-1α is a suggested dual-function cytokine that diverged from IL-1β in mammals potentially by acquiring additional biological roles that relate to highly conserved regions in the pro-domain of IL-1α, including a nuclear localisation sequence (NLS) and histone acetyl transferase (HAT)-binding domains. Why evolution modified pro-IL-1α’s subcellular location and protein interactome, and how this shaped IL-1α’s intracellular role, is unknown. TurboID proximity labelling with pro-IL-1α suggested a nuclear role for pro-IL-1α that involved interaction with HATs, including EP300. We also identified and validated inactivating mutations in the pro-IL-1α NLS of multiple mammalian species. However, HAT-binding domains were also conserved in species that had lost pro-IL-1α nuclear localisation. Together, these data suggest that HAT binding and nuclear localisation occurred together, and that while some species lost the NLS in their pro-IL-1α, HAT binding was maintained. The NLS was lost from several distinct species at different evolutionary times, suggesting convergent evolution, and that the loss of the NLS confers some important biological outcome.

## Introduction

Inflammation is typically a beneficial, coordinated response of immune cells that protects a host from pathogenic infection. Communication between immune cells is driven by signalling molecules called cytokines, which are usually secreted proteins that bind to receptors on target cells to elicit a signalling response. However, there are examples where some cytokines appear to have dual function through an additional intracellular role (*1*). One of the most important cytokine families in the host response to infection is the interleukin-1 (IL-1) family, which consists of 11 members of both pro- and anti-inflammatory cytokines. Pro-inflammatory members of the IL-1 family include the well-studied cytokines IL-1α and IL-1β (*2*). IL-1β is an ancient protein conserved throughout vertebrates, while IL-1α arose as a gene duplication of IL-1β and is present only in mammals (*3*).

Like IL-1β, IL-1α is produced as a less active 31 kDa precursor (pro-IL-1α), that requires enzymatic processing to a fully active form, and signals through the type 1 IL-1 receptor when released. However, the pro-domains of IL-1α and IL-1β appear to have evolved separate biological roles (*3*). The pro-domain of IL-1β in mammals exhibits limited sequence conservation, and it is considered simply to stop the activation and secretion of IL-1β (*4*). The pro-domain of IL-1α, on the other hand, is highly conserved amongst mammals and has several domains which may have evolved through a functional specialisation, driving the divergence of IL-1α from IL-1β (*3*). Highly conserved domains within the IL-1α pro-domain include a nuclear localisation sequence (NLS), a motif of basic amino acids that facilitates trafficking to the nucleus, and histone acetyl transferase (HAT)-binding domains (*3*). HATs (also known as lysine acetyl transferases (KATs)) can be located in the nucleus or cytosol, and they mediate the acetylation of lysine residues on proteins, neutralising their positive charge and regulating the function of the protein (*5*). Although originally identified for their role in histone modification and chromatin remodelling, HATs have since been characterised for their specialised role in DNA-damage repair, regulating gene expression and the modification of non-histone substrates (*6*). Previously published studies have identified a possible role for pro-IL-1α in the regulation of gene expression via binding to HAT complexes (*7-9*), although the exact nature of its nuclear role remains unclear.

Here we aimed to discover the intracellular function of IL-1α by establishing a pro-IL-1α interactome using proximity labelling and by analysing evolutionary evidence. By tagging pro-IL-1α with the enhanced biotin ligase mutant, TurboID (*10*), we were able to efficiently biotinylate nearby and interacting proteins of pro-IL-1α in cells and establish a pro-IL-1α interactome, revealing a network of new IL-1α interactors including several HATs. We validated that the toothed whale clade and the rodent suborder castorimorpha, along with several other mammalian species, have lost NLS-dependent nuclear localisation of pro-IL-1α, but maintained HAT-binding domain conservation. Our data suggest the most likely role of intracellular pro-IL-1α is to influence HAT activity, and highlights that IL-1α subcellular location has been manipulated by evolutionary pressures on several independent occasions.

## Results

### Characterisation of pro-IL-1α proximity labelling

We previously established the highly conserved nature of the pro-domain of pro-IL-1α in mammals in comparison to pro-IL-1β, suggesting additional functions of the pro-domain of IL-1α other than regulating proteolytic cleavage (*3*). The pro-domain of pro-IL-1α contains an NLS, which facilitates nuclear import (*11*). The majority of mammals have a highly conserved NLS in their pro-domain of pro-IL-1α, suggesting a possible nuclear role. Therefore, we investigated the nuclear interactors of pro-IL-1α by using TurboID proximity labelling to define a pro-IL-1α interactome (*10*). HeLa cells were transiently transfected to express either human pro-IL-1α, pro-IL-1α-TurboID, or TurboID alone (Figure 1A). We then characterised the sub-cellular distribution and expression levels of the constructs using immunocytochemistry. Pro-IL-1α was largely localised to the nucleus (Figure 1B) consistent with previous reports of ectopically and endogenously expressed IL-1α (*12, 13*). The expression pattern of pro-IL-1α-TurboID closely matched that of untagged pro-IL-1α, confirming that the TurboID tag did not affect expression or distribution of the protein (Figure 1B). When expressed alone, TurboID was diffusely present throughout the entire cell (Figure 1B). Western blotting was also used to validate protein expression and labelling (Figure 1C). Addition of biotin (500µM, 30min) to transfected cells followed by streptavidin-HRP labelling of either fixed cells, or of protein gels, confirmed the protein expression patterns described above (Figure 1B) and that proteins within the cell were biotinylated (Figure 1D). Treatment of pro-IL-1α- and pro-IL-1α-TurboID-expressing cells with the calcium ionophore ionomycin, an established activator of calpain-dependent pro-IL-1α processing (*14*), confirmed that the TurboID tag did not interfere with IL-1α processing, or the kinetics of IL-1α processing, suggesting normal trafficking and processing (Figure 1E). These data confirm that pro-IL-1α-TurboID and TurboID alone were expressed correctly, trafficked normally, and that the biotin ligase activity of the TurboID was functional.

**Figure 1.**
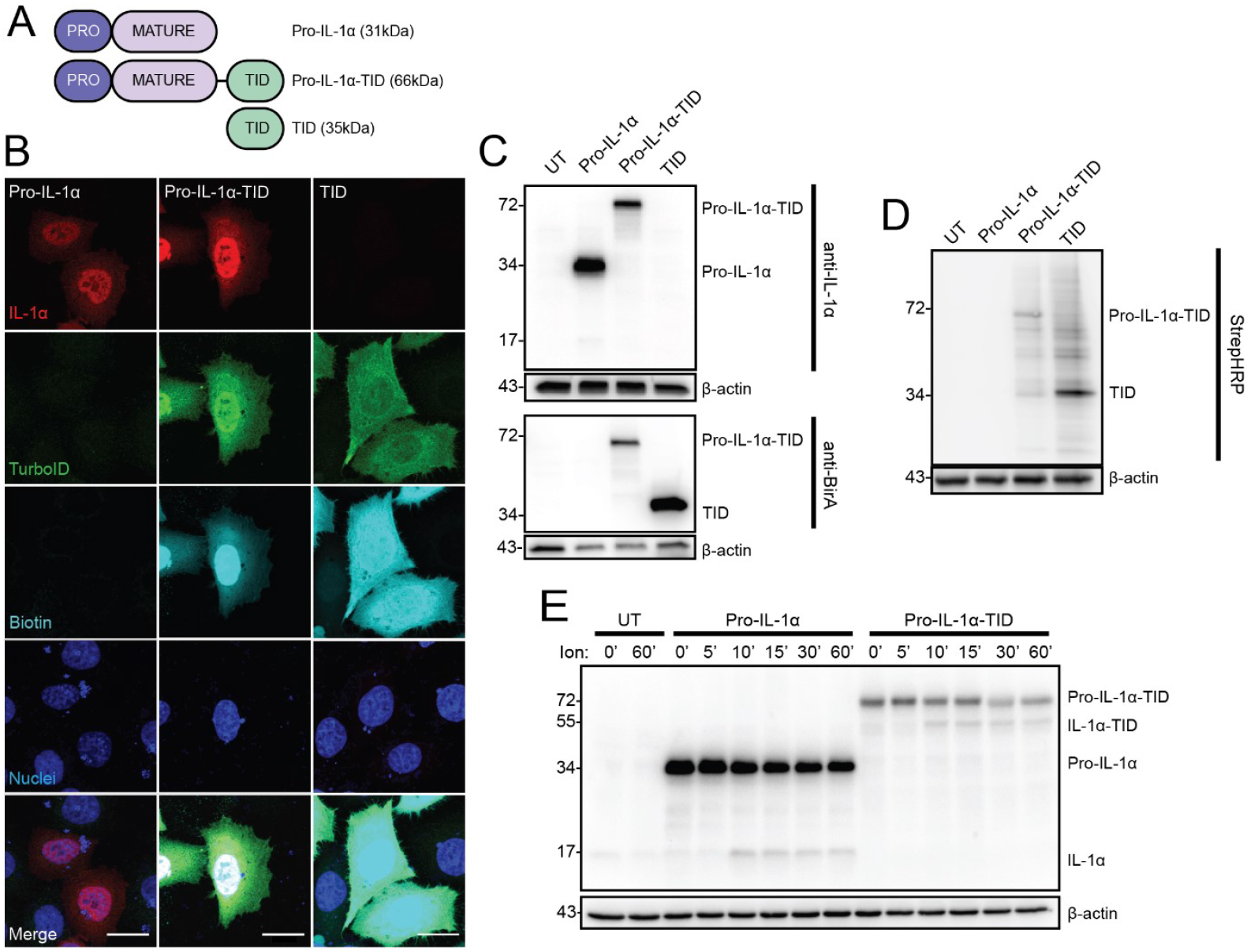
Characterisation of pro-IL-1α-TurboID in HeLa cells. (**A**) Schematic of pro-IL-1α, pro-IL-1α-TurboID (pro-IL-1α-TID) and TurboID (TID) constructs. (**B**) HeLa cells were transfected with constructs in (A), then treated with biotin (500 µM, 30 min), and analysed by immunofluorescence microscopy (n=2). Anti-IL-1α labels pro-IL-1α and mature IL-1α, anti-BirA labels TurboID, streptavidin-HRP labels biotinylated proteins. Dark blue represents nuclei stained by DAPI. Scale bars are 10 µm. (**C, D**) HeLa cells were untransfected (UT) or transfected with constructs in (A), then treated with biotin (500 µM, 30 min). Cell lysates were probed for **(C)** IL-1α (anti-IL-1α), TurboID (anti-BirA) (n=4), and **(D)** biotin (StrepHRP) (n=4). (**E**) HeLa cells were untransfected (UT) or transfected with pro-IL-1α or pro-IL-1α-TID and were then treated with ionomycin (Ion) for 0, 5, 10, 15, 30 or 60 minutes (n=4). Cell lysates were probed for IL-1α by western blotting. 0 and 60 minutes were used as representatives for UT cells.

### Pro-IL-1α interactome highlights a prominent proximity to HATs

To establish a pro-IL-1α interactome, HeLa cells transfected with TurboID alone or pro-IL-1α-TurboID were treated with biotin (500 µM, 30 min), and samples were analysed for biotinylated proteins by mass spectrometry (Figure 2A). Principal component analysis established that the pro-IL-1α-TurboID interactome was different from the TurboID alone interactome (Figure 2B). Comparison of pro-IL-1α-TurboID and TurboID alone highlighted 56 selectively enriched proteins within the pro-IL-1α interactome (Figure 2C; Supplementary Table 1). Subcellular location analysis using ingenuity pathway analysis (IPA) of significantly enriched proteins indicated that proteins interacting with pro-IL-1α-TurboID were largely associated with the nucleus (Figure 2D; Supplementary Figure 1). Since only a few nuclear proteins were significantly enriched in the pro-IL-1α-TurboID interactome, and because some nuclear proteins were significantly enriched in TurboID alone, this suggested that the pro-IL-1α interacting proteins identified were specific to pro-IL-1α itself (Supplementary Figure 2). Further subcellular location analysis of significantly enriched nuclear proteins revealed an enrichment at chromosomes (Figure 2E). IPA was also used to determine the canonical pathways and biological functions predicted to be associated with the pro-IL-1α interactome based on the representation of the 56 significant proteins within these processes (Figure 2F). Multiple nuclear receptor signalling pathways were significantly enriched, including the aryl hydrocarbon receptor, glucocorticoid receptor, vitamin D receptor, thyroid receptor and retinoic acid receptor signalling pathways. Further to this, gene expression was the most significant biological function associated with the pro-IL-1α interactome (Figure 2G). Based on a database of the current literature, STRING (https://string-db.org/) analysis determined known interacting partners within the pro-IL-1α interactome and used Gene Ontology (GO) terms to identify functions of the pro-IL-1α interactome proteins. This analysis identified many members of the pro-IL-1α interactome as HAT complex proteins (Figure 3). The interaction between pro-IL-1α and the HAT protein, EP300 was previously identified (*7, 13*). Other HAT proteins identified here that have not previously been recorded to interact with pro-IL-1α are CRT2C, NCOA2, NCOA3, ZZZ3, SUPT20H, EPC1, DMAP1, YEATS2 and EP400 (Figure 3). These data highlight an association between pro-IL-1α and proteins that regulate gene expression. More specifically, the identification of many HAT-interacting proteins within the pro-IL-1α interactome is a strong indication of an intracellular role of pro-IL-1α.

**Figure 2.**
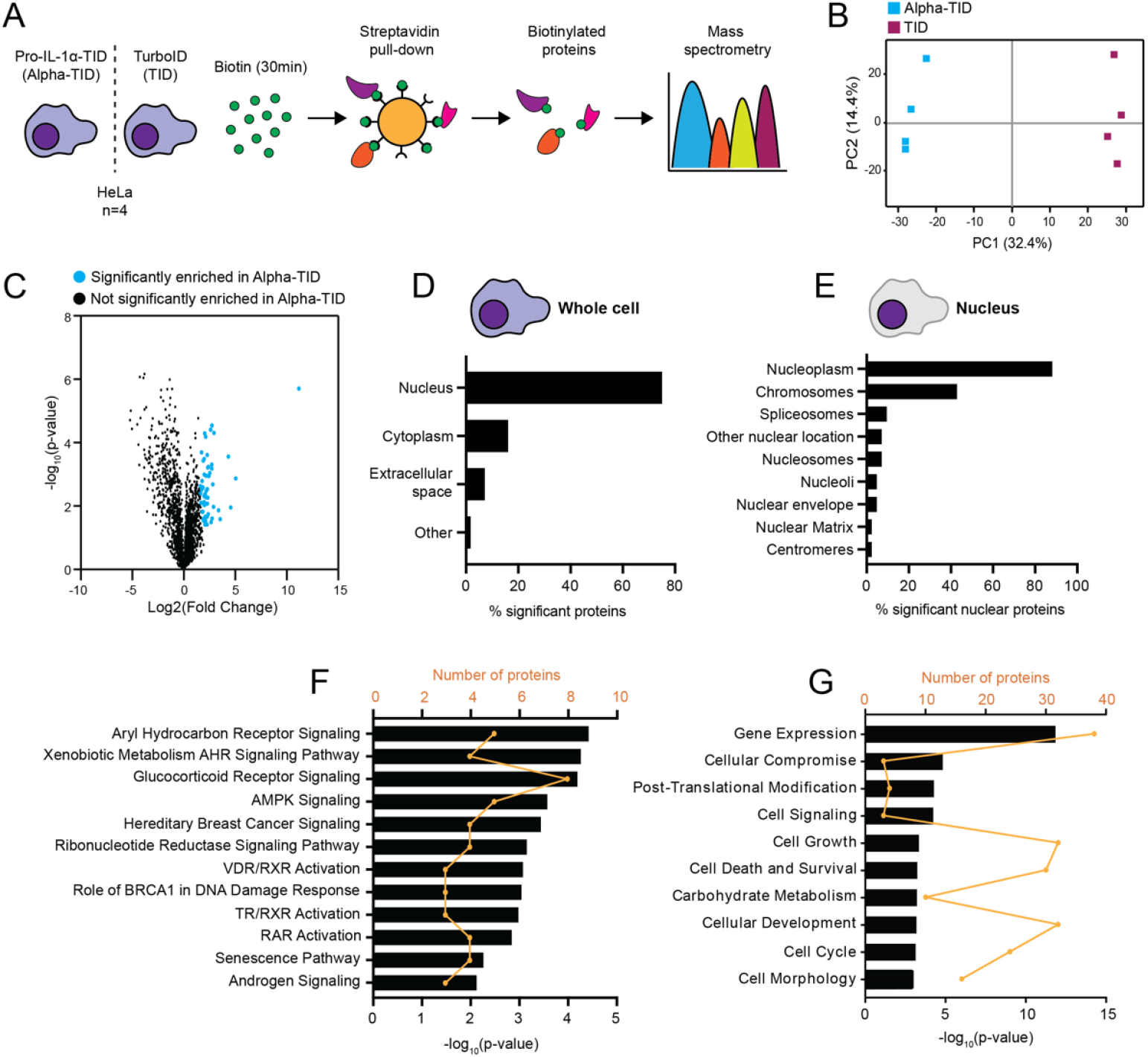
Spatial and functional profiling of pro-IL-1α-TurboID. (**A**) Schematic illustrating the pipeline of biotinylation enrichment analysis by mass spectrometry. HeLa cells were transfected with pro-IL-1α-TurboID (Alpha-TID) or TurboID (TID), and then treated with biotin (500 µM, 30 min, n=4). Streptavidin pull-down of biotinylated proteins was performed, and biotinylated proteins were analysed by mass spectrometry. (**B**) Principal component analysis of LFQ intensities. A single square denotes an independent experiment. Experimental groups (Alpha-TID or TID) are grouped by colour. (**C**) Volcano plot of proteins enriched in Alpha-TID (blue) compared to TID following two-sample t-test of LFQ intensity values (significance determined as s0=2; FDR=0.01). (**D-G**) Ingenuity pathway analysis (IPA) of proteins significantly enriched in Alpha-TID. (**D**) Subcellular location of all significantly enriched proteins. (**E**) Nuclear location of significantly enriched nuclear proteins. (**F**) Canonical pathways (p<0.01) and (**G**) biological functions (top 10) associated with proteins significantly enriched in Alpha-TID. Black bars represent log^10^(p-value) of overlap between Alpha-TID proteins and the proteins within each canonical pathway or biological function and were obtained via the right-tailed Fisher’s exact test implemented in the IPA software. Orange line denotes number of Alpha-TID proteins present within each canonical pathway or biological function.

**Figure 3.**
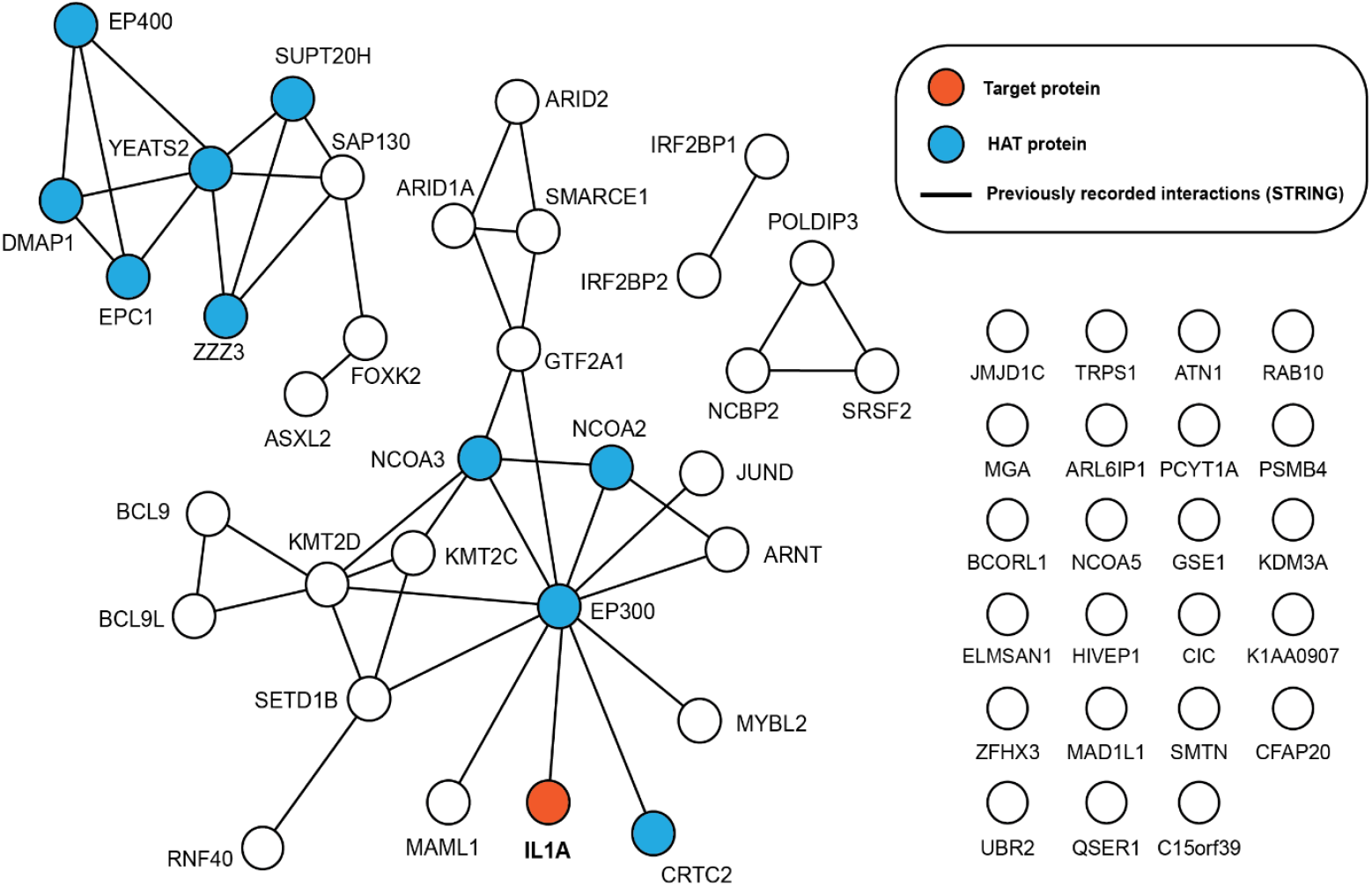
STRING analysis of pro-IL-1α-TurboID biotinylated proteins highlighting prominent proximity to HAT proteins. STRING network analysis (generated via https://string-db.org/) of proteins significantly enriched in pro-IL-1α-TurboID. Lines represent previously recorded physical interactions identified by STRING, unconnected proteins have no previously recorded interactions with significantly enriched proteins. Histone acetyltransferase (HAT) proteins were identified through gene ontology (GO) analysis. Proteins are displayed using gene symbol label. Gene symbol for the target protein (IL1A; pro-IL-1α) represents the target protein of this experiment.

### Several mammalian species, including all castorimorpha, lack pro-IL-1α NLS conservation

Although the NLS is well conserved in the pro-IL-1α of most mammals (Figure 4A), we recently identified that in the toothed whale clade the pro-IL-1α NLS contained mutations in the ‘KKRR’ motif, resulting in predicted loss of NLS function (*3*). Given recent advances in sequence availability, we were able to investigate if pro-IL-1α contained a non-functional NLS in other mammals. We retrieved and aligned 226 pro-IL-1α amino acid sequences from 157 mammalian species, and assessed the conservation of the pro-IL-1α NLS KKRR motif in these species. We identified numerous mammalian species, in addition to toothed whales, which contain amino acid mutations in the pro-IL-1α NLS KKRR motif leading to a predicted loss of NLS function (Figure 4B, Supplementary Figure 3A). We also identified the KKRR motif nucleotide sequences for these species (Supplementary Figure 3B). These species were the naked mole-rat, southern grasshopper mouse, American beaver, Ord’s kangaroo rat, banner-tailed kangaroo rat, Chinese pangolin, big brown bat, Tasmanian devil, koala and common brushtail. There may have been additional mammalian species that also have NLS mutations for which we did not retrieve an IL-1α sequence. We subsequently analysed the divergence times of these species, which suggested that the pro-IL-1α NLS loss events we detected were evolutionary distinct, as they occurred in diverse mammalian lineages (Figure 4B) (*15*). These data suggested that multiple mammalian species may have lost pro-IL-1α nuclear localisation following distinct evolutionary events.

**Figure 4.**
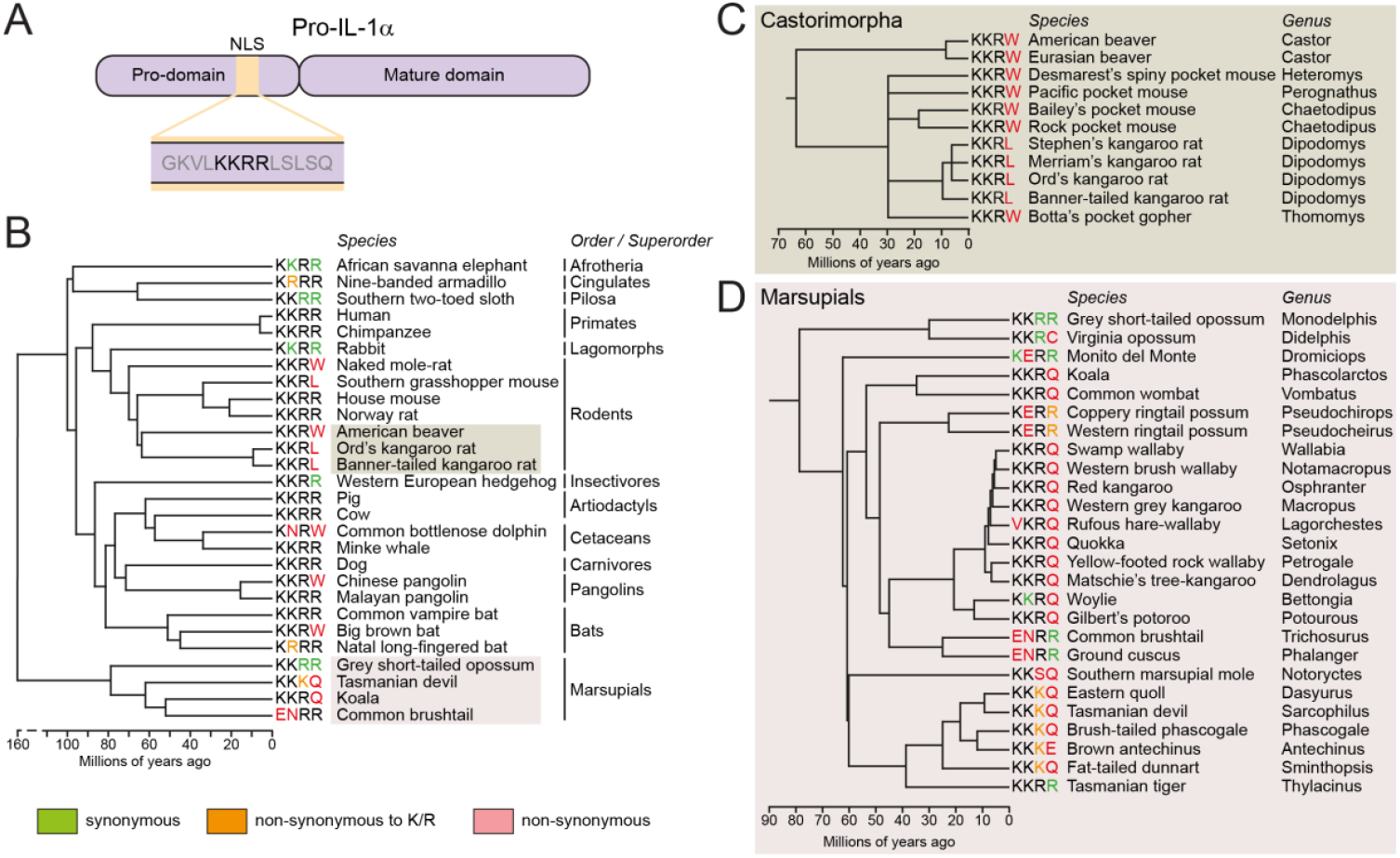
Several mammalian species lack pro-IL-1α NLS conservation. (**A**) Schematic of pro-IL-1α protein domains, including the NLS location and amino acid sequence. (**B**) Model mammalian evolutionary tree, highlighting representative mammalian species from various orders and superorders, as well as the identified species with NLS mutations in the KKRR motif, including (**C**) an expanded model tree of the rodent suborder castorimorpha, and (**D**) an expanded model tree of the marsupial lineage, only including species for which an IL-1α sequence could be retrieved. Non-synonymous mutations are shown in red, apart from mutations to K/R, which are shown in orange. Divergence times were retrieved using TimeTree (15).

To more precisely identify the evolutionary time-point when the NLS mutations arose in these lineages, and given that the initial alignment performed did not include sequences from all mammalian species, where possible we retrieved IL-1α exon 3 (which contains the NLS coding sequence) DNA or RNA sequence reads from species closely related to the identified NLS mutant species (Supplementary Figure 4A). This approach also allowed us to retrieve IL-1α nucleotide sequences from species whose genome or RNA had been sequenced but not necessarily annotated for IL-1α. We then compared whether the KKRR motif nucleotide sequence region from these reads matched that of the human KKRR motif nucleotide sequence, which was conserved across many mammalian species (Supplementary Figure 3B). Sequence reads from the NLS mutant species themselves were also assessed to validate the presence of the mutated KKRR motif sequences. We identified that, similar to the toothed whale clade, all species of the rodent suborder castorimorpha, which contains beavers, pocket mice, kangaroo rats and gophers, for which we could obtain sequence reads possessed non-synonymous nucleotide mutations in the KKRR motif, resulting in an amino acid substitution of the final arginine residue to tryptophan or leucine (Figure 4C, Supplementary Figure 4B, C). These data suggested that the NLS mutation arose after the castorimorpha lineage diverged from its rodent ancestors approximately 67 million years ago, and was subsequently conserved. Most of the castorimorpha species possessed a KKRW motif, but the kangaroo rat sequences analysed contained a KKRL motif, caused by an additional non-synonymous nucleotide mutation in the final amino acid residue that occurred in kangaroo rats after they further diverged (Figure 4C, Supplementary Figure 4B). Through this sequence read analysis, we were able to validate the loss of the NLS in the naked mole-rat, southern grasshopper mouse, Chinese pangolin and big brown bat, but the close relatives of these species that we assessed all possessed an intact NLS, with no apparent non-synonymous nucleotide mutations in the KKRR motifs (Supplementary Figure 5-8). Thus, the NLS loss was specific to these individual mutant NLS species, and these were isolated, separate evolutionary events. The marsupial lineage contained multiple species with varying non-synonymous nucleotide mutations in the NLS KKRR motif, indicating an almost global lack of NLS conservation in this mammalian group (Figure 4D, Supplementary Figure 9, 10).

### KKRR motif mutations result in reduced pro-IL-1α nuclear localisation

We had identified species that possessed mutations in the pro-IL-1α NLS sequence, resulting in predicted loss of pro-IL-1α nuclear localisation, but we needed to validate this experimentally. Therefore, we transfected HeLa cells with either full length human pro-IL-1α, full length human pro-IL-1β which does not have an NLS and therefore is localised to the cytosol, a chimeric IL-1 protein composed of a human IL-1α pro-domain fused to the human IL-1β mature domain (pro-α-mat-β), or a chimeric IL-1α protein composed of a toothed whale (*Orcinus orca*, killer whale) IL-1α pro-domain fused to the human IL-1α mature domain (Orca pro-IL-1α) (Figure 5A), and then assessed cytokine subcellular localisation. Full length pro-IL-1α was predominantly located in the nucleus of the cells, whereas full length pro-IL-1β was largely cytosolic (Figure 5B, C). Fusing the IL-1α pro-domain to the IL-1β mature domain resulted in nuclear localisation of this protein, confirming that nuclear localisation is an inherent phenotype driven by the IL-1α pro-domain (Figure 5B, C). The chimeric Orca pro-IL-1α was distributed throughout the cell cytoplasm, confirming that the Orca pro-domain of pro-IL-1α does not contain a functional NLS, and therefore that toothed whale pro-IL-1α is not trafficked to the nucleus, but instead is predominantly located in the cytosol (Figure 5B, C).

**Figure 5.**
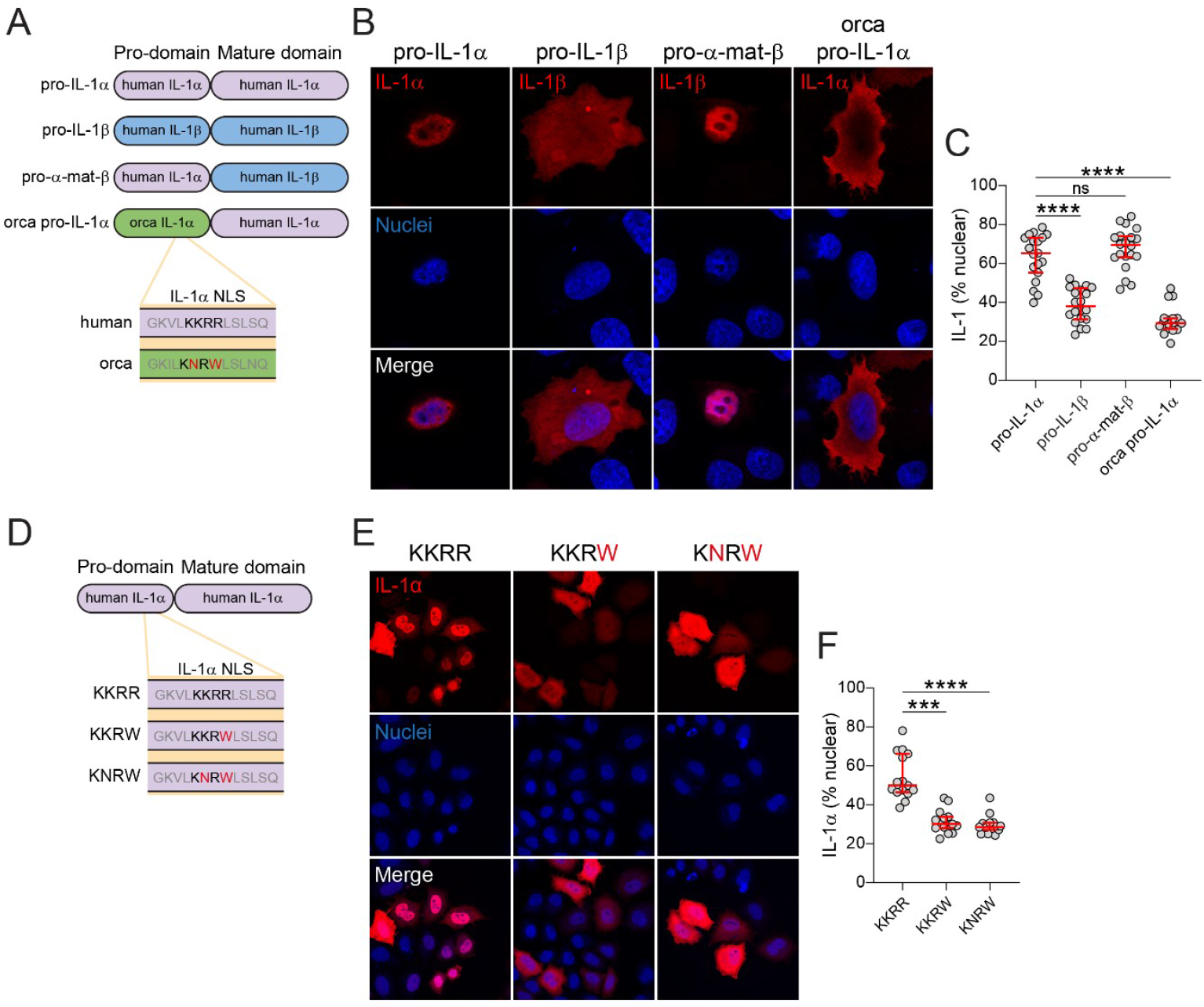
KKRR motif mutations in the NLS cause reduced pro-IL-1α nuclear localisation. (**A-C**) HeLa cells were transfected with full length human pro-IL-1α, full length human pro-IL-1β, human IL-1α pro-domain fused to the human IL-1β mature domain (pro-α-mat-β), or Orca IL-1α pro-domain fused to the human IL-1α mature domain (Orca pro-IL-1α). (**A**) Schematic of constructs. (**B**) Representative immunofluorescence images and (**C**) quantification of IL-1 nuclear localisation are shown (n=19-21 fields of view from four independent experiments). (**D-F**) HeLa cells were transfected with human pro-IL-1α plasmids containing a KKRR, KKRW, or KNRW NLS motif. (**D**) Schematic of constructs. (**E**) Representative maximum projection confocal immunofluorescence images and (**F**) quantification of IL-1α nuclear localisation are shown (n=15 fields of view from three independent experiments). Data are median ± IQR. Data were analysed using one-way ANOVA followed by Dunnett’s post-hoc test (C) or Kruskal-Wallis test followed by Dunn’s post-hoc test (F). ***P<0.001; ****P<0.0001; ns, not significant.

It was possible that the reduction in nuclear localisation caused by the Orca IL-1α pro-domain may have been due to differences in the rest of the pro-IL-1α NLS sequence, and not specifically due to the mutations in the KKRR motif that resulted in a KNRW motif instead. To determine whether mutations in the KKRR motif alone were sufficient to cause a reduction in nuclear localisation, we transfected HeLa cells with either human WT pro-IL-1α (KKRR) or a mutated human pro-IL-1α containing the toothed whale KNRW motif within its NLS (Figure 5D). WT pro-IL-1α was strongly localised to the nucleus, whereas introduction of the double amino acid mutation (KNRW) from toothed whales into the pro-IL-1α construct reduced pro-IL-1α nuclear localisation, and enhanced its cytosolic distribution, indicating that these mutations alone were sufficient for loss of NLS function (Figure 5E, F). Whilst the toothed whale NLS sequences typically contained two amino acid mutations in the KKRR motif, resulting in KNRW, the newly identified NLS mutant species typically only contained a mutation in the final arginine residue of this motif, most commonly to KKRW, which incidentally is the same sequence as the sperm whale, another member of the toothed whale clade (Figure 5D). Thus, to establish whether a single amino acid mutation in the KKRR motif was sufficient to cause a reduction in pro-IL-1α nuclear localisation, we also transfected HeLa cells with human pro-IL-1α containing a KKRW motif within its NLS, and this too reduced pro-IL-1α nuclear localisation, indicating loss of NLS function (Figure 5E, F). These data suggest that the loss of KKRR motif conservation in the newly identified NLS mutant species, such as castorimorpha and marsupials, would result in reductions in pro-IL-1α nuclear localisation, similar to toothed whales.

### HAT-binding domains are conserved in most mammalian species, including pro-IL-1α NLS mutants

Binding motifs for HATs have previously been described in the pro-domain of pro-IL1α, and HATs have been described to interact with pro-IL-1α (*7*). Having validated the loss of pro-IL-1α NLS function in numerous mammalian species, this meant we could further explore the evolutionary relationship between pro-IL-1α nuclear localisation and HAT binding. To do this, we assessed the conservation of these HAT-binding domains in the mammalian species with an intact NLS and compared this to species with a mutated KKRR motif sequence. Conservation of the HAT-binding domains appeared consistent between the intact and mutant NLS species (Figure 6A, B). The N-terminal HAT-binding domain (amino acid residues 7-19) was much better conserved in comparison to the other HAT-binding domain (amino acid residues 98-108) (Figure 6B). There was no clear difference in conservation of the whole pro-domain between species with an intact NLS and NLS mutants (Figure 6C). Marsupials had moderate conservation of the N-terminal HAT-binding domain, and poor conservation of the second, although these species had poor conservation across the whole IL-1α pro-domain (Figure 6A-C) and mature domain compared to other species (Supplementary Figure 11). This poor sequence conservation in marsupials may also explain why pro-IL-1α NLS mutations were detected in most marsupial species (Figure 1B), and why marsupials have poor conservation of the thrombin cleavage site in IL-1α (*16*). Thus, the species in which the NLS was mutated retained similar conservation of HAT-binding domains in the absence of an intact pro-IL-1α NLS, potentially suggesting that the intracellular role of pro-IL-1α was driven by HAT binding, and not by nuclear localisation solely due to the NLS. This suggests that the function of the pro-IL-1α NLS remains unclear and may have evolved to facilitate interactions with nuclear HATs, or for another reason entirely.

**Figure 6.**
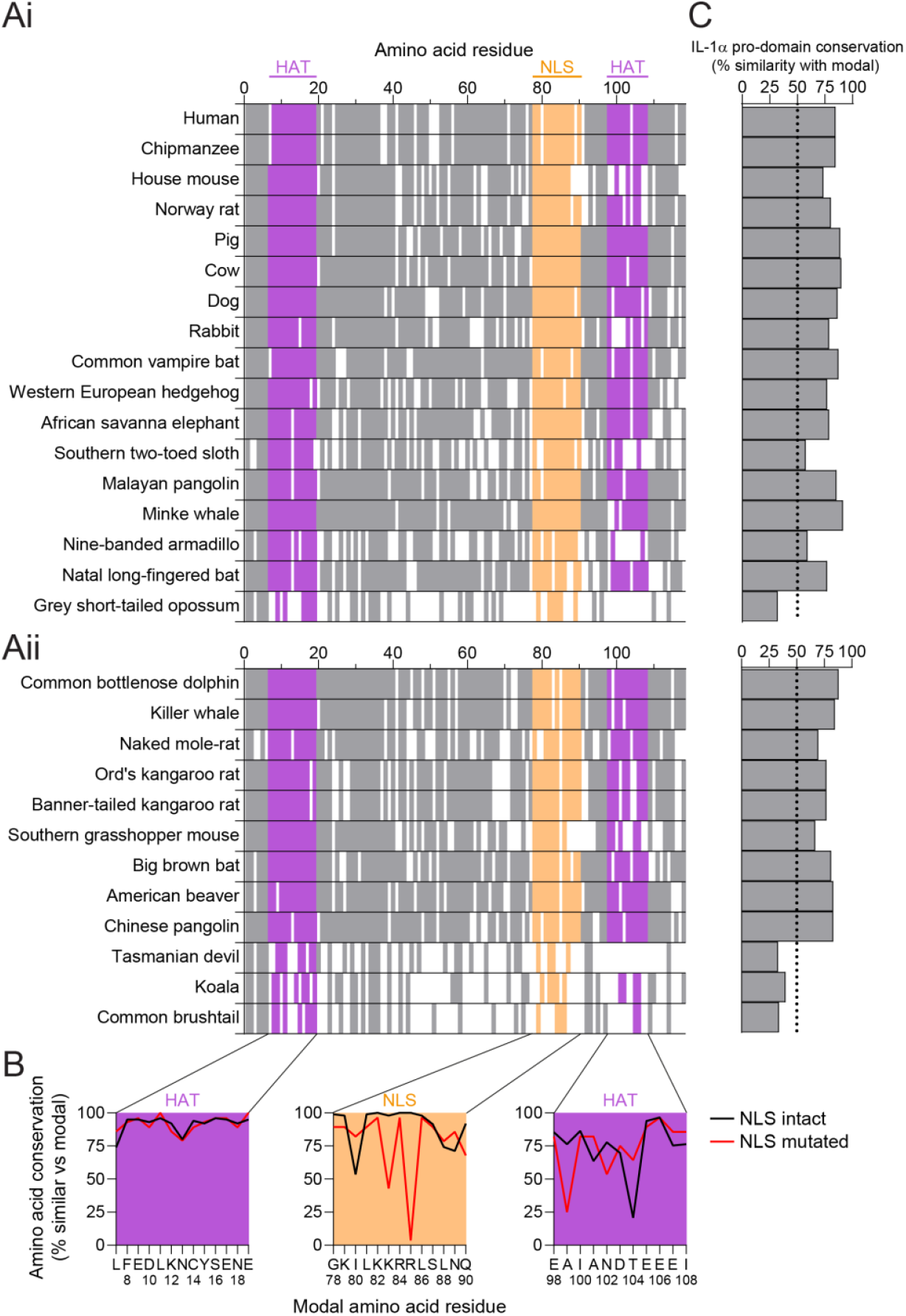
HAT-binding domains are similarly conserved in most mammalian species, including NLS mutants. (**A**) Pro-IL-1α pro-domain amino acid conservation in representative intact NLS (Ai) and mutated NLS (Aii) species. Residues that match the modal amino acid from the full sequence alignment are indicated by a solid colour, whereas residues that did not match are indicated by a white gap. Positions where gaps were the modal residue were removed from the alignment. HAT-binding domains are identified in purple (amino acids 7-19, 98-108 in human sequence), and the NLS is identified in orange (amino acids 78-90 in human sequence). (**B**) Amino acid sequence conservation of both HAT-binding domains and the NLS for all sequences with an intact or mutated NLS (relative to the modal amino acid residue from the full alignment of all sequences). (**C**) Conservation of the whole pro-IL-1α pro-domain relative to the modal pro-domain sequence.

### Monotremes have an intact NLS in pro-IL-1α, and evidence of HAT-binding domain conservation

To further dissect the temporal origin of the NLS and HAT-binding regions, we analysed the IL-1α gene region in monotremes, which diverged from the mammalian lineage approximately 180-185 million years ago (20-25 million years before marsupials), and whose genomes have recently been sequenced (*17*) (Figure 7A). We located the IL-1α gene in the platypus and short-beaked echidna genome assemblies, although this was unannotated in the platypus, and observed similar chromosomal anatomy to other mammals, although the *CENPT* gene was located between *IL1B* and *IL1A* in both monotreme species (Figure 7B). We next determined the IL-1α amino acid and nucleotide sequences for these species by locating the pro-IL-1α exons, and identified a relatively well conserved NLS sequence that was predicted to be functional, and that had a fully conserved KKRR motif (Figure 7C, D, Supplementary Figure 12). Despite this conserved NLS, both monotreme pro-IL-1α sequences were relatively poorly conserved compared to modal amino acid sequence from all the mammalian species analysed, similar to the marsupials (Figure 7E, F). It was unsurprising that monotreme and marsupial IL-1α was considerably different to placental mammals, as these mammalian groups diverged from placental mammals more than 150 million years ago, and it is possible that the marsupial and monotreme sequences may more closely resemble the ancestral IL-1α protein.

**Figure 7.**
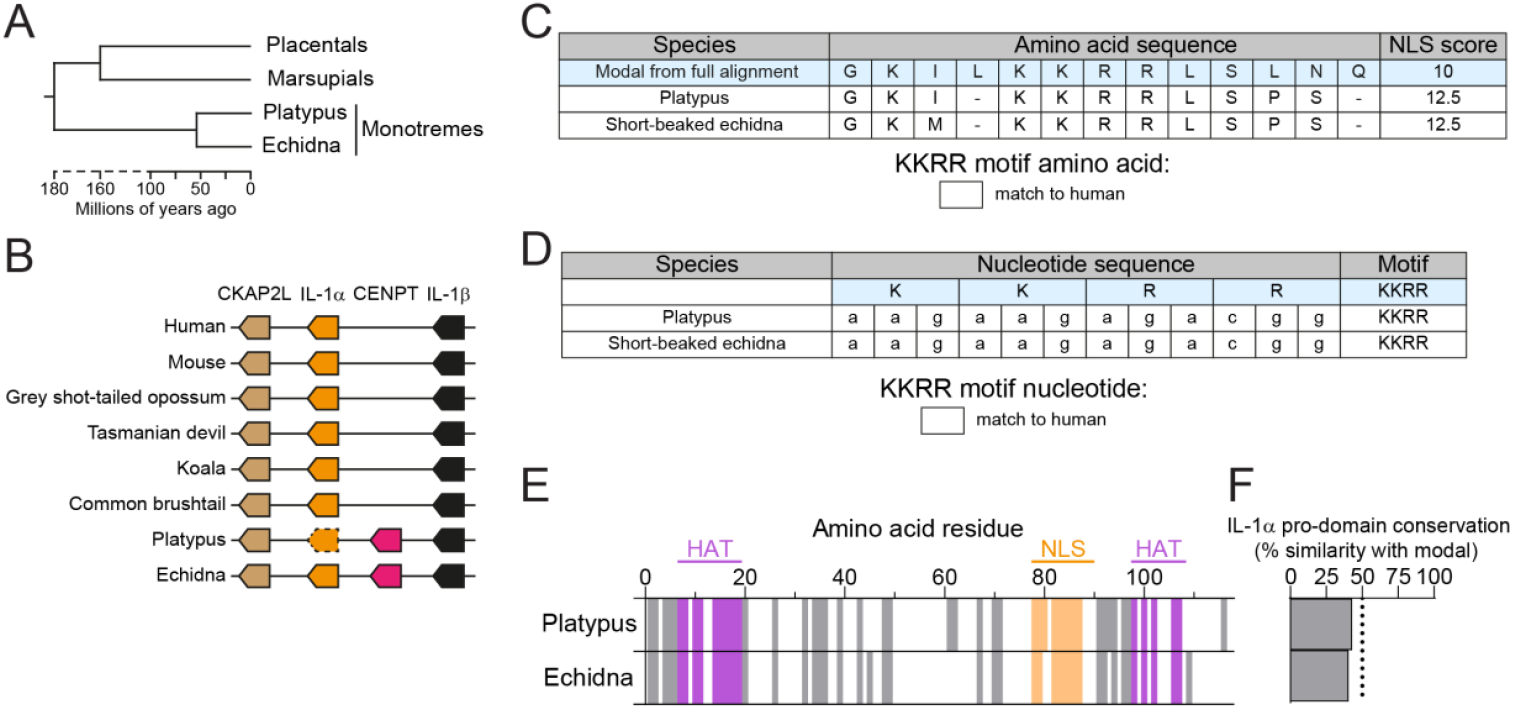
Monotremes have an intact NLS in pro-IL-1α, and evidence of HAT-binding domains. (**A**) Model evolutionary tree of the divergence of placental mammals, marsupials and monotremes. Divergence times were retrieved using TimeTree (15). (**B**) Chromosomal anatomy of IL-1α gene region in several mammalian species. Arrow direction indicates gene orientation. Dotted outline indicates lack of genome annotation. Gene lengths and distances between genes are not to scale. (**C**) Pro-IL-1α NLS amino acid sequences of the platypus and short-beaked echidna. Modal sequence was taken from the alignment of IL-1α from all isoforms of all 158 species. NLS scores were estimated using NLSmapper. Sequences correspond to G^78^-Q^90^ in human sequence. (**D**) KKRR motif nucleotide sequences for the two monotreme species. (**E**) Pro-IL-1α pro-domain amino acid conservation in the platypus and short-beaked echidna. Residues that match the modal amino acid from the full sequence alignment are indicated by a solid colour, whereas residues that did not match are indicated by a white gap. Positions where gaps were the modal residue were removed from the alignment. HAT-binding domains are identified in purple (amino acids 7-19, 98-108 in human sequence), and the NLS is identified in orange (amino acids 78-90 in human sequence). (**F**) Conservation of the whole pro-IL-1α pro-domain relative to the modal pro-domain sequence.

We then examined the HAT-binding regions of monotreme pro-IL-1α to see if they were also conserved. The N-terminal HAT-binding domain was moderately conserved in comparison to the other HAT-binding domain, which was poorly conserved (Figure 7E). We also identified regions in the IL-1α pro-domain (amino acids 1-6 and 32-36) that were highly conserved across monotremes, marsupials and placental mammals, suggesting that these regions may be of fundamental importance for pro-IL-1α function. Thus, these data would place the evolution of the NLS and possibly the N-terminal HAT-binding domain after the IL-1 gene duplication event in the proto-mammal but before mammalian divergence, driving IL-1α’s divergence from IL-1β, and the NLS was subsequently lost in certain mammals in distinct evolutionary events. However, from these data we are unable to conclude whether the NLS or HAT-binding domain appeared first.

## Discussion

IL-1α is a pro-inflammatory cytokine that has been considered to act as a damage-associated molecular pattern (DAMP), or an alarmin, upon its release from damaged cells, and it is implicated in the inflammation contributing to many conditions, including respiratory, cardiovascular, and neurological diseases (*18, 19*). IL-1α signalling is also suggested to contribute to other inflammatory processes, including cellular senescence where it promotes the senescence-associated secretory phenotype (SASP) (e.g. (*20, 21*)). However, pro-IL-1α is a putative dual-function cytokine, as it is also characterised by a nuclear sub-cellular localisation and potential function (*1*). Nuclear localisation of pro-IL-1α is driven by an NLS within its pro-domain (*11, 12*). Our lab previously reported that, whilst the NLS within the pro-domain of pro-IL-1α is conserved amongst many mammalian sequences, it is lost in the toothed whale clade (*3*). We previously used this to test the hypothesis that nuclear functionalisation drove the evolution of IL-1α (*3*). We predicted that if the NLS was important, then there would be an increase in the amino acid replacements in the pro-domain of toothed whale IL-1α, as without an NLS, the nuclear function would now be redundant. However, the sequence of the pro-domain of toothed whale IL-1α remains highly conserved, despite the functional mature domain of IL-1α acquiring an increase of amino acid replacements. This suggests that a nuclear functionalisation related specifically to the NLS was probably not a major force driving the divergence of IL-1α (*3*). However, as we present in this study, even in pro-IL-1α sequences from species where there is no functional NLS, there are conserved HAT-binding domains. This, and the interactome described above, suggest that HAT binding may be the primary intracellular role of pro-IL-1α, and that nuclear trafficking driven by the NLS served some auxillary role. This purpose of the NLS may be simply to facilitate binding to HATs that reside in the nucleus, or it may be completely independent of HAT binding, and instead related to the trafficking and release of IL-1α, although this requires further investigation.

We identified nine histone-modifying enzymes as part of the pro-IL-1α interactome, in addition to the HAT protein, EP300, which is reported to interact with pro-IL-1α in the nucleus (*7, 9, 13*). Histone modification is a critical process in the regulation of gene expression, which is a key biological function associated with the pro-IL-1α interactome identified in this study. This suggests that pro-IL-1α may play an important role in regulating gene expression via its association with proteins involved in histone modification, further supporting its role as a dual functioning protein.

The mutations causing a loss of the NLS arose at distinct, independent time points in mammalian evolution. Castorimorpha lost NLS function after divergence from the rodent lineage approximately 67 million years ago (*15, 22, 23*), which would place this shortly before the cretaceous-paleogene mass extinction event that resulted in the loss of 75% of all species, including non-avian dinosaurs. The precise timings of mammalian diversification around this extinction event are unclear (*24*), however, with the divergence of castorimorpha also estimated between 50-55 million years ago (*25*). In toothed whales, the loss of the NLS occurred approximately 35 million years ago when they diverged from baleen whales (*15*). The loss of KKRR motif conservation and predicted NLS function in marsupials may be related to the geographical isolation of Australasian and South American marsupials upon separation of the supercontinent Gondwana, with Australasian marsupials such as the koala and Tasmanian devil losing KKRR motif conservation, whereas the South American opossums retained it. The monito del monte, although an inhabitant of South America, has been taxonomically classified as a closely related sister group of Australasian marsupials that likely diverged shortly before these species migrated to Australia, possibly explaining why this species has also lost predicted NLS function (*26*).

The evolutionary convergence on loss of NLS function, not just in toothed whales but in the rodent suborder castorimorpha, marsupials and other mammalian lineages, suggests that modifying the subcellular location of pro-IL-1α conferred an evolutionary advantage to these species. However, the events that drove selection for NLS loss are unclear, as are the benefits conferred by this mutation, and they may be due to reducing the nuclear function of IL-1α. It is also possible that the NLS serves as a mechanism to regulate the secretion of IL-1α by limiting available pro-IL-1α for processing and secretion by virtue of its sub-cellular compartmentalisation (*13, 27*). Thus, a reduction of IL-1α nuclear localisation may enhance its extracellular release. Better understanding the implications of losing IL-1α nuclear localisation in these species may reveal valuable insights into IL-1α biology.

Critically, in all these independent mutation events in which NLS functionality was lost, at least one of the pro-IL-1α HAT-binding domains remained highly conserved, suggesting that HAT binding is still applying purifying selection pressure. This could be explained by the NLS mutations preventing active trafficking of pro-IL-1α into the nucleus, but not preventing its passive diffusion. Thus, the strongly reduced, but not completely absent, nuclear localisation of pro-IL-1α may have still been sufficient to retain HAT-binding conservation in the mutant NLS species. Another possibility is that cytosolic pro-IL-1α can interact with cytosolic HATs (or KATs) in the mutant NLS species. The functions of cytosolic HATs can be independent of their acetylation activity, for example the multifunctional EP300 HAT protein is also reported to exhibit E3 and E4 ubiquitin ligase activity (*28*). Interestingly, this ligase activity is only functional when EP300 is in the cytosolic fraction of the cell (*29*). Pro-IL-1α has been identified to be poly-ubiquitinated and this promotes its processing into the inflammatory mature form, but the ligases involved in the process have not yet been identified (*30*). Here we have shown that pro-IL-1α interacts with EP300 and at least two other E3 ligases proteins (RNF40 and UBR2, Supplementary Table 1). Nevertheless, while cytosolic interactions of pro-IL-1α and HAT enzymes may be important, the vast majority of mammalian species have a highly conserved NLS and HAT-binding domains. Furthermore, we have shown here that HAT enzymes and other gene regulatory proteins such as transcription factors are the most enriched group of proteins in the interactome of pro-IL-1α, suggesting nuclear HAT interaction to be the most likely deciding factor in pro-IL-1α evolution.

Thus, IL-1α is likely to be a dual-function cytokine, with an intracellular role dictated by a conserved pro-domain and HAT binding, and an extracellular pro-inflammatory role executed by the mature domain. The loss of the NLS from different mammalian species at distinct evolutionary times, suggesting convergent evolution, indicates an important role for pro-IL-1α subcellular location. Understanding the intracellular role of pro-IL-1α, its interactions with HATs and the consequences of this to a cell and organism may represent the next challenge in understanding IL-1α.

## Materials and Methods

### Reagents

Goat anti-human IL-1α (AF-200-NA) was from R&D Systems. Rabbit anti-human BirA (11582-RP01) was from Stratech Scientific. Bovine serum albumin (A9647), Anti-β-actin-peroxidase (A3854) and biotin (B4639) were from Sigma-Aldrich. Lipofectamine 3000, Streptavidin-HRP conjugate (S911), Alexa Fluor™ 488 donkey anti-goat IgG (A-11055), Alexa Fluor™ 594 donkey anti-goat IgG (A-11058), Streptavidin Alexa Fluor™ 594 conjugate (S11227) or Alexa Fluor 647™ donkey anti-rabbit IgG (A-31573), DAPI (D1306) and ProLong Gold antifade mounting reagent (P36934) were from Invitrogen. Cytiva Amersham ECL Prime Western Blotting Detection Reagent (RPN2236) was from GE Healthcare. Protease inhibitor cocktail (539131) was from Merck Millipore. Fugene HD (E2311) was from Promega. Rabbit anti-goat IgG (P044901-2) or goat anti-rabbit IgG (P044801-2) secondary antibodies were from Agilent.

### Cell culture

HeLa cells were cultured in Dulbecco’s modified Eagle’s medium (DMEM) containing 10% foetal bovine serum (FBS) and 100 U ml^-1^ penicillin and 100 µg ml^-1^ streptomycin, and were incubated at 37 °C and 5% CO^2^. Before experiments, cells were seeded at a densities of 5 × 10^4^ cells ml^-1^ or 1 × 10^5^ cells ml^-1^ in 24-well plates for immunocytochemistry experiments, or 2 × 10^5^ cells ml^-1^ in 24-well or 10 cm culture dishes for western blotting and proximity labelling experiments, respectively, and left overnight.

### Plasmids

Coding sequences were obtained from human *IL1A* (NCBI Gene ID: 3552), human *IL1B* (NCBI Gene ID: 3553) and *Orcinus orca IL1A* (NCBI reference sequence: XM_004276766.2). Pro-IL-1α, pro-α-mat-β, orca-pro-(h)IL-1α and pro-IL-1β genes were synthesized and cloned to pcDNA3.1^(+)^ vectors (Life Technologies). pro-α-mat-β chimeric expressed the Met^1^to Arg^112^ of human IL-1α, followed by Tyr^113^ to Ser^269^ of human IL-1β. Orca-pro-(h)IL-1α chimeric expressed the Met^1^ to Ile^108^ of Orca IL-1α, followed by Ile^109^ to Ala^271^ of human IL-1α. To generate IL1a lentiviral expression vectors, Addgene plasmid #50919 was digested with BstBI and BstXI and a synthesised fragment comprising a multiple cloning site (MCS) and bGH polyA was inserted via restriction cloning, to create pLV-Ef1a-MCS. This plasmid was digested with KpnI and NotI and used as the backbone for the following constructs. hIL1a WT (NLS coding sequence aagaagagacgg); pro-IL-1α was amplified from pcDNA3.1-pro-IL-1α, using F1F and F2R. The NLS mutant (KKRW;aagaagagaTgg); was generated via a two fragment HiFi assembly using primers to introduce the mutation. Fragment 1, F1F and aF1R, Fragment 2, aF2F and F2R. The second NLS mutant (KNRW; aagaaCagaTgg); was again generated by two fragment HiFi assembly using primers F1F and bF1R for fragment 1, and bF2F and F2R for fragment 2. To generate a pro-IL-1α-TurboID expression vector, the pro-IL-1α sequence was amplified from pcDNA3.1-pro-IL-1α using pF1F and pF1R, and TurboID sequence amplified from Addgene plasmid #107173 using primers tF2F and tF2R. The vector was assembled by HiFi assembly (NEB) into pcDNA3.1/KpnI/NotI. To generate the TID expression vector, the TurboID sequence was amplified from Addgene plasmid #107173 using primers tF1F and tF1R. Followed by HiFi assembly into pLV-Ef1a-MCS/KpnI/NheI.

### Transient transfections

For experiments using the chimeric IL-1 constructs, HeLa cells were transfected for 18-24 h with 250 ng of human pro-IL-1α, human pro-IL-1β, human pro-α-mat-β, or Orca pro-IL-1α using Lipofectamine 3000, according to manufacturer’s instructions. For experiments using the single or double amino acid pro-IL-1α constructs, HeLa cells were transfected for 24 h with 250 ng of full length human WT pro-IL-1α, or pro-IL-1α containing the KKRW or KNRW mutations using FuGENE HD transfection reagent at a ratio of 3:1 according to manufacturer’s instructions. For TurboID experiments, HeLa cells were transfected for 18 h with either pro-IL-1α (500 ng), pro-IL-1α-TurboID (500 ng) or TurboID alone (12.5 ng) per 5 × 10^5^ cells seeded using Lipofectamine 3000. For mass spectrometry analysis, after transfection of cells with pro-IL-1α-TurboID or TurboID alone, cell media was changed to complete DMEM and left to incubate for a further 24 h prior to stimulation to obtain an optimal cell number for stimulation and cell scraping.

### Immunocytochemistry

Cells were washed once in PBS then fixed in 4% paraformaldehyde (PFA) for 20 min before being permeabilized with PBS, 0.1% Triton X-100 (PBST) for 5 min at room temperature. Cells were incubated in blocking solution (5% BSA, PBST) for 1 h at room temperature, before incubation with goat anti-human IL-1α (400 ng ml^-1^), rabbit anti-human BirA (1:500 v/v) or streptavidin-HRP (1 µg ml^-1^) primary antibodies in 1% BSA PBST at 4 °C overnight. Cells were then incubated with Alexa Fluor™ 488 donkey anti-goat IgG (4 µg ml^-1^), Alexa Fluor™ 594 donkey anti-goat IgG (4 µg ml^-1^), Alexa Fluor 647™ donkey anti-rabbit IgG (4 µg ml^-1^) or Streptavidin Alexa Fluor™ 594 conjugate (2 µg ml^-1^) secondary antibodies for 1 h at room temperature. Nuclei were stained with DAPI (0.5 µg ml^-1^). Coverslips were mounted on slides using ProLong Gold antifade mounting reagent and left to dry overnight.

### Microscopy and nuclear localisation analysis

Widefield microscopy images were acquired for experiments using the chimeric IL-1 constructs using a Zeiss Axioimager.D2 upright microscope using a 20×/0.5 EC Plan-neofluar objective and captured using a Coolsnap HQ2 camera (Photometrics) through Micromanager software v1.4.23. Specific band pass filter sets for DAPI, FITC and Texas red were used to prevent bleed through from one channel to the next. Images were acquired from 3-5 fields of view from each independent experiment, and nuclear localisation was quantified using the ImageJ ‘Intensity Ratio Nuclei Cytoplam’ tool and expressed as percentage of total IL-1 fluorescence intensity that co-localised with DAPI signal.

Confocal microscopy images were acquired for experiments using the single or double amino acid pro-IL-1α constructs, or for TurboID experiments, using a 63×/1.40 HCS PL Apo objective on a Leica TCS SP8 AOBS inverted or upright confocal microscope with LAS X software (v3.5.1.18803). To prevent interference between channels, lasers were excited sequentially for each channel. The blue diode with 405 nm and the white light laser with 488nm, 594 nm and 647 nm laser lines were used, with hybrid and photon-multiplying tube detectors with detection mirror settings set appropriately. Z-stacks were acquired with 0.3 µm steps between Z sections. Images were acquired from 5 fields of view from each independent experiment, and nuclear localisation was quantified manually using maximum projections and expressed as the percentage of total IL-1α fluorescence intensity that co-localised with DAPI signal.

### Biotinylation experiments

HeLa cells transfected with pro-IL-1α-TurboID or TurboID alone were treated with biotin (500 µM) for 30 min in serum-free DMEM. Cells were then washed 4 times in cold PBS (containing CaCl^2^ and MgCl_2_), scraped and pelleted at 1000 x g for 5 min at 4 °C.

### MaxQuant analysis, filtering, and transformation

Streptavidin pull-down and mass spectrometry analysis was commercially outsourced and carried out by Sanford Burnham Prebys Proteomics Core, La Jolla. Mass spectrometry was carried out on dried and isolated biotin-enriched peptides following biotinylation in pro-IL-1α-TurboID or TurboID groups. Mass spectrometry data were analysed by MaxQuant and quantification was provided as LFQ (label-free quantification) intensities for both peptides and protein groups. LFQ intensities for protein groups were used as the unit for investigation from this point forward. The Perseus computational platform (Tyanova et al., 2016; https://maxquant.net/perseus/) was used to prepare Protein group LFQ intensities for pro-IL-1α-TurboID and TurboID. Protein groups were loaded into Perseus and LFQ intensities were identified as the columns of interest in this analysis. The dataset was then filtered to remove potential contaminants, reverse hits and proteins only identified by site. Data were transformed using a log2(x) transformation and normal distribution was checked by plotting histograms of LFQ intensity for all repeats within each experimental group (pro-IL-1α-TurboID or TurboID alone). Samples were then grouped according to biological replicates and rows were filtered so that at least one experimental group (pro-IL-1α-TurboID or TurboID alone) contained a valid value for all four replicates. Imputation was carried out to allow any further missing values to be replaced with values predicted from a normal distribution. Principal component analysis (PCA) was performed on each biological replicate within pro-IL-1α-TurboID and TurboID alone groups.

### Ingenuity pathway analysis

Ingenuity’s pathway analysis (IPA, Qiagen) was used to analyse pathway enrichment in pro-IL-1α-TurboID significantly enriched proteins. Subcellular location analysis was achieved by IPA using gene ontology (GO) database of subcellular location terms. For canonical pathway and biological function analysis, the p-value of overlap was calculated according to the overlap between pro-IL-1α-TurboID significantly enriched proteins and proteins present with each pathway/function.

### STRING analysis

Gene names for proteins significantly enriched in pro-IL-1α-TurboID were uploaded into STRING (https://string-db.org/). Network shows “physical subnetwork” meaning all physical interactions previously reported in the literature and functional annotations were exported containing all gene ontology terms associated with the pro-IL-1α-TurboID interactome. HAT proteins were determined by filtering gene ontology terms for “containing: histone, and containing: acetyltransferase”.

### Western blotting

Western blot analysis was performed on HeLa cell lysates for IL-1α, BirA and streptavidin. Supernatant was removed and cells were lysed in lysis buffer (50 mM Tris-HCl, 150 mM NaCl; Triton-X-100 1% v/v, pH 7.3) containing protease inhibitor cocktail. Equal volumes of lysates were run on 12% SDS-polyacrylamide gels and transferred at 25 V onto PVDF membranes using a Trans-Blot^®^ Turbo Transfer™ System (Bio-Rad). Membranes were blocked in 5% BSA (w/v) in PBS, 0.1% Tween (v/v) for 1 h at room temperature. Membranes were incubated at 4°C overnight with goat anti-human IL-1α (200 ng ml^-1^), rabbit anti-human BirA (1:1000 v/v) and Streptavidin-HRP (500 ng ml^-1^) in 1% BSA (w/v) in PBS, 0.1% Tween. Membranes were washed in PBS Tween and incubated in rabbit anti-goat IgG (500 ng ml^-1^) or goat anti-rabbit IgG (250 ng ml^-1^) in 1% BSA (w/v) in PBS, 0.1% Tween for 1 h at room temperature. Proteins were visualised with Cytiva Amersham ECL Prime Western Blotting Detection Reagent and G:BOX (Syngene) and Genesys software. Membranes were then probed for β-actin as a loading control.

### Protein alignment

IL-1α amino acid sequences were retrieved from the NCBI orthologues database (Ref_Seq) (accessed 05/08/2021). In total, 226 amino acid sequences from 157 species were obtained, including all available isoforms from each species. Sequences were aligned using MUSCLE on MEGAX using default settings and the NLS sequences (^78^GKVLKKRRLSLSQ^90^ in human) were analysed. HAT-binding domain conservation was assessed by comparing the aligned IL-1α pro-domain sequences to the modal amino acid sequence from the full alignment. The HAT-binding domains previously identified in pro-IL-1α (amino acids 7-19 and 98-108 in human pro-IL-1α) were compared between species with a functional and non-functional NLS.

### NLS score

NLS score was calculated using NLS mapper (*31*) using the full IL-1α amino acid sequence, or part of the exon 3 amino acid sequence obtained from the SRA analysis below, selecting the highest monopartite NLS score containing the KKRR motif region.

### SRA analysis

We obtained DNA and RNA sequencing read data for the mutant NLS species, as well as closely related species, from the NCBI sequence read archive database (https://www.ncbi.nlm.nih.gov/sra). We then used BLAST (default parameters, optimised for megablast or blastn) to align these sequence reads against an IL-1α exon 3 nucleotide query sequence (obtained from NCBI genome viewer) of the respective closely related NLS mutant species, to identify reads that closely matched the query sequence. Where possible, up to three sequence reads were obtained from different animals, from different studies. We then converted up to the top 100 sequence reads that had an alignment score >80 to a multiple sequence alignment using the NCBI multiple sequence alignment viewer. For each nucleotide position coding for the KKRR motif in the IL-1α NLS (aagaagagacgg in human), the % of bases in the sequence reads that matched the human sequence base was calculated.

### Data presentation and analysis

Nuclear localisation quantification from immunofluorescence images is presented as median ± interquartile range, with each data point representing a field of view, and was assessed for normal distribution using Shapiro-Wilk normality test and analysed using one-way ANOVA with Dunnett’s post-hoc test, or Kruskal-Wallis test with Dunn’s post-hoc test for non-parametric data. TurboID experiments were analysed as stated in the methods. Western blots are representative of 4 independent experiments. Statistical analysis was performed using GraphPad Prism (v9).

## Acknowledgements

D.B. and C.H. were funded by Medical Research Council (MRC) grant MR/T016515/1. R.W. was funded by BHF Accelerator Award AA/18/4/34221. V.S.T. was funded by CONICYT (Becas Chile 721704488). G.L.-C. was funded by Medical Research Council (MRC) grant MR/T016043/1. The Bioimaging Facility microscopes used in this study were purchased with grants from BBSRC, Wellcome and the University of Manchester Strategic Fund

## Author Contributions

Conceptualisation: D.B., G.L.-C. and C.H.

Methodology: H.B. and A.A.

Investigation: R.W., V.T., and C.H.

Visualisation: R.W. and C.H.

Funding acquisition: D.B. and G.L.-C.

Supervision: D.B., G.L.-C., J.P.G. and C.H.

Writing - original draft: R.W., C.H., and D.B.

Writing - review and editing: R.W., C.H., D.B., J.P.G., V.T., G.L.-C., J.R.-A., H.B. and A.A.

## Data availability

The data the support the findings of this study are available from the corresponding author upon reasonable request.

