## Supplementary data for "Proximity labelling and evolutionary evidence reveal insights into IL-1α nuclear networks and function"

### Supplementary Materials

**Supplementary Table 1. Pro-IL-1α-TurboID significantly enriched proteins.** 56 proteins significantly enriched via pro-IL-1α-TurboID-mediated biotinylation. Significance calculated using Perseus computational platform statistical analysis (s0=2 and FDR=0.01). Table shows gene name, protein name, log2 fold change (FC) and -log10 (p-value).

| Gene name | Protein name | Log2 FC | -Log10 p-value |
|---|---|---|---|
| IL1A | Interleukin-1 alpha | 11.16 | 5.67 |
| EP300 | Histone acetyltransferase p300 | 2.72 | 4.54 |
| YEATS2 | YEATS domain-containing protein 2 | 2.60 | 4.41 |
| CRTC2 | CREB-regulated transcription coactivator 2 | 2.92 | 4.30 |
| DMAP1 | DNA methyltransferase 1-associated protein 1 | 2.03 | 4.30 |
| ZFHX3 | Zinc finger homeobox protein 3 | 2.12 | 4.18 |
| ARID2 | AT-rich interactive domain-containing protein 2 | 1.72 | 3.69 |
| KMT2D | Histone-lysine N-methyltransferase 2D | 2.26 | 3.61 |
| ZZZ3 | ZZ-type zinc finger-containing protein 3 | 4.32 | 3.56 |
| SMTN | Smoothelin | 1.96 | 3.48 |
| QSER1 | Glutamine and serine-rich protein 1 | 1.89 | 3.43 |
| TRPS1 | Zinc finger transcription factor Trps1 | 2.71 | 3.31 |
| MYBL2 | Myb-related protein B | 2.39 | 3.24 |
| JMJD1C | Probable JmjC domain-containing histone demethylation protein 2C | 2.32 | 3.22 |
| NCOA3 | Nuclear receptor coactivator 3 | 2.71 | 3.18 |
| MAML1 | Mastermind-like protein 1 | 2.51 | 3.04 |
| CIC | Protein capicua homolog | 1.99 | 3.00 |
| SUPT20H | Transcription factor SPT20 homolog | 2.17 | 2.95 |
| ARNT | Aryl hydrocarbon receptor nuclear translocator | 2.43 | 2.94 |
| KMT2C | Histone-lysine N-methyltransferase 2C | 5.03 | 2.88 |
| BCORL1 | BCL-6 corepressor-like protein 1 | 1.77 | 2.79 |
| SMARCE1 | SWI/SNF-related matrix-associated actin-dependent regulator of chromatin subfamily E member 1 | 2.80 | 2.68 |
| ATN1 | Atrophin-1 | 1.60 | 2.63 |
| KDM3A | Lysine-specific demethylase 3A | 1.81 | 2.59 |
| BCL9L | B-cell CLL/lymphoma 9-like protein | 1.86 | 2.58 |
| HIVEP1 | Zinc finger protein 40 | 2.27 | 2.53 |

| | | | |
|---|---|---|---|
| ARID1A | AT-rich interactive domain-containing protein 1A | 1.73 | 2.52 |
| NCOA5 | Nuclear receptor coactivator 5 | 2.18 | 2.48 |
| EP400 | E1A-binding protein p400 | 1.63 | 2.40 |
| C15orf39 | Uncharacterized protein C15orf39 | 2.07 | 2.38 |
| ASXL2 | Putative Polycomb group protein ASXL2 | 1.97 | 2.32 |
| EPC1 | Enhancer of polycomb homolog 1 | 2.13 | 2.28 |
| BCL9 | B-cell CLL/lymphoma 9 protein | 2.32 | 2.27 |
| ELMSAN1 | ELM2 and SANT domain-containing protein 1 | 1.97 | 2.12 |
| UBR2 | E3 ubiquitin-protein ligase UBR2 | 1.66 | 2.11 |
| MAD1L1 | Mitotic spindle assembly checkpoint protein MAD1 | 1.91 | 2.08 |
| POLDIP3 | Polymerase delta-interacting protein 3 | 2.28 | 2.08 |
| IRF2BP1 | Interferon regulatory factor 2-binding protein 1 | 2.03 | 2.05 |
| CFAP20 | Cilia- and flagella-associated protein 20 | 1.98 | 2.03 |
| SAP130 | Histone deacetylase complex subunit SAP130 | 2.88 | 1.97 |
| SETD1B | Histone-lysine N-methyltransferase SETD1B | 4.55 | 1.96 |
| JUND | Transcription factor jun-D | 3.37 | 1.87 |
| SRSF2 | Serine/arginine-rich splicing factor 2 | 1.84 | 1.82 |
| KIAA0907 | UPF0469 protein KIAA0907 | 1.80 | 1.81 |
| GSE1 | Genetic suppressor element 1 | 2.36 | 1.75 |
| NCOA2 | Nuclear receptor coactivator 2 | 2.27 | 1.67 |
| FOXK2 | Forkhead box protein K2 | 2.08 | 1.67 |
| ARL6IP1 | ADP-ribosylation factor-like protein 6-interacting protein 1 | 1.98 | 1.62 |
| IRF2BP2 | Interferon regulatory factor 2-binding protein 2 | 2.73 | 1.61 |
| RAB10 | Ras-related protein Rab-10 | 3.53 | 1.58 |
| PSMB4 | Proteasome subunit beta type-4 | 1.89 | 1.58 |
| PCYT1A | Choline-phosphate cytidylyltransferase A | 1.90 | 1.54 |
| RNF40 | E3 ubiquitin-protein ligase BRE1B | 2.11 | 1.53 |
| GTF2A1 | Transcription initiation factor IIA subunit 1 | 2.64 | 1.49 |
| NCBP2 | Nuclear cap-binding protein subunit 2 | 2.25 | 1.42 |
| MGA | MAX gene-associated protein | 2.06 | 1.41 |

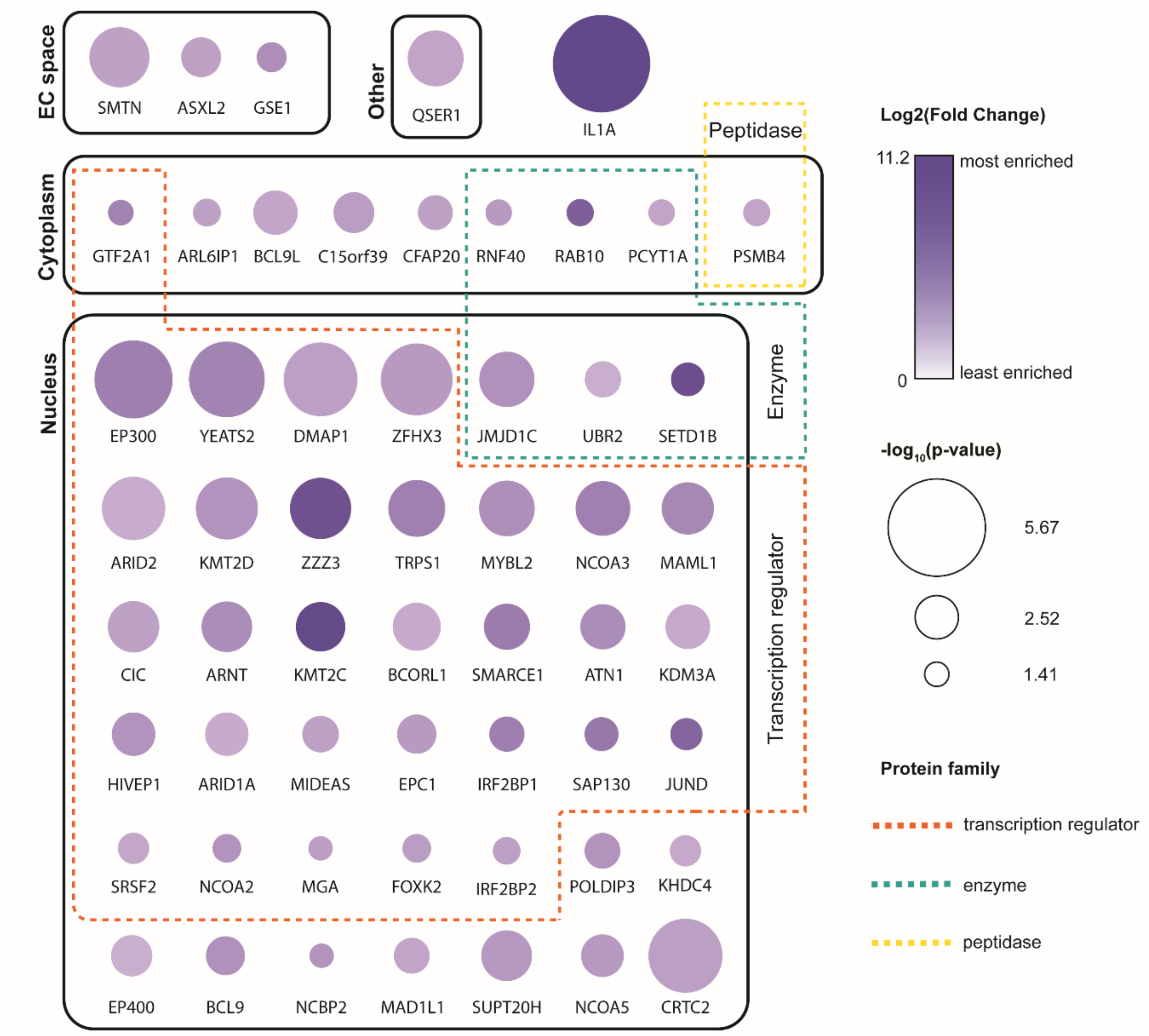


**Supplementary Figure 1. Subcellular location and protein family groups of pro-IL-1α-TurboID biotinylated proteins.** Proteins are clustered by subcellular location and protein family as described by IPA. Those proteins with no assigned protein family group were categorised as “other” by IPA. Gene symbol name is used to annotate each protein. Proteins are coloured by fold change in enrichment, displayed as Log2 (Fold Change), and sized by p-value, displayed as $-\log_{10}$ (p-value), determined by two-sample t-test of LFQ intensity values (significance determined as s0=2; FDR=0.05). Fold change and p-value are calculated following comparison with TID control. Subcellular locations are determined by IPA and grouped by solid boxes (EC space: extracellular space). Protein families are grouped by dashed lines; transcription regulator (orange); enzyme (green); peptidase (yellow).

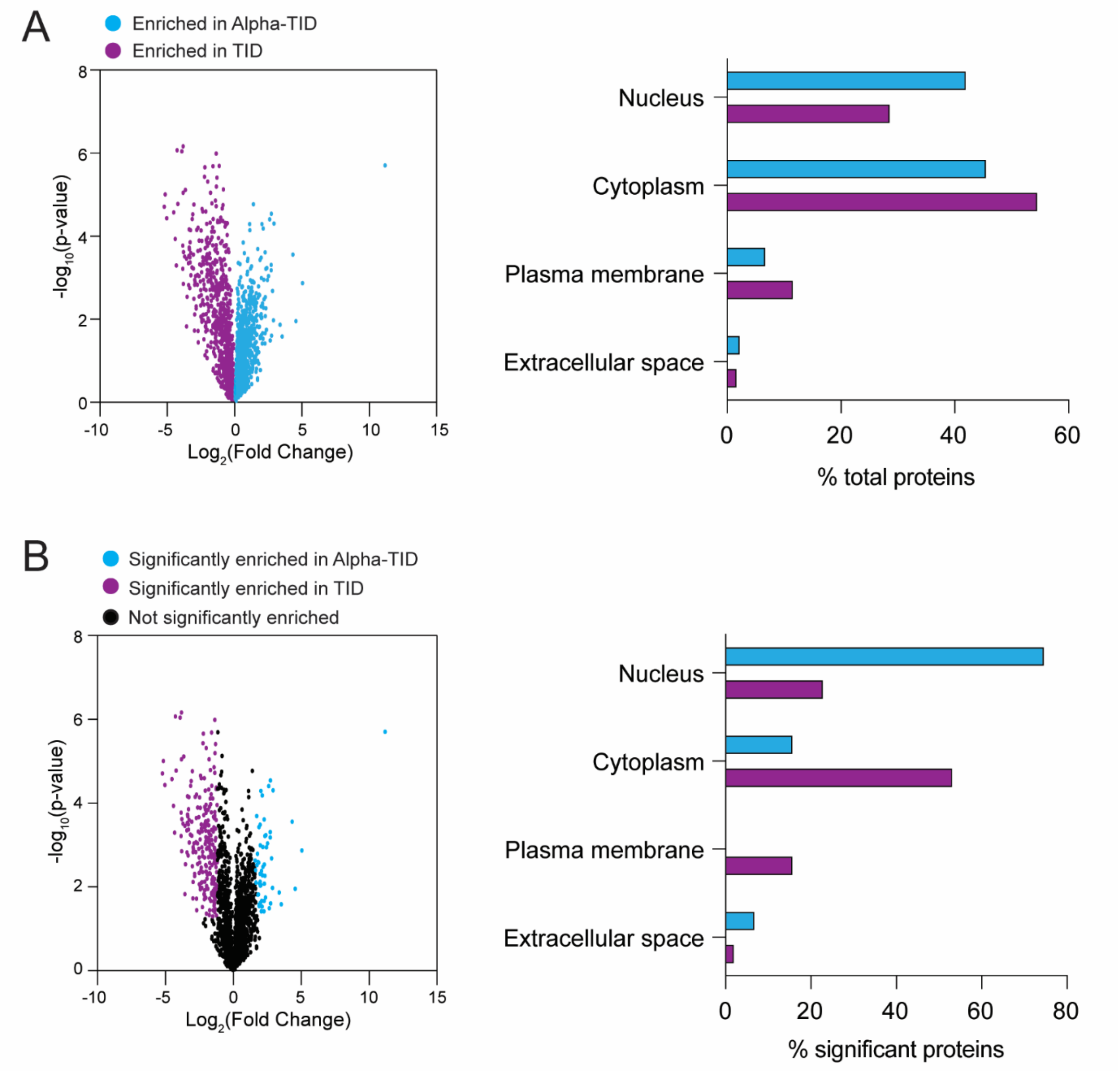


**Supplementary Figure 2. Subcellular location of pro-IL-1α-TurboID and TurboID enriched proteins.** (**A**) Volcano plot highlighting Alpha-TID and TID enriched proteins. Subcellular location of Alpha-TID and TID enriched proteins. (**B**) Volcano plot highlighting Alpha-TID and TID significantly enriched proteins. Subcellular location of Alpha-TID and TID significantly enriched proteins.

A

| Species | Amino acid sequence | | | | | | | | | | | | | | NLS score |
|---|---|---|---|---|---|---|---|---|---|---|---|---|---|---|---|
| *Intact NLS* | | | | | | | | | | | | | | | |
| Human | G | K | V | L | K | K | R | R | L | S | L | - | S | Q | 9.5 |
| Chipmanzee | G | K | V | L | K | K | R | R | L | S | L | - | S | Q | 9.5 |
| House mouse | G | K | I | L | K | K | R | R | L | S | F | - | S | E | 10.5 |
| Norway rat | G | K | I | L | K | K | R | R | L | S | F | - | N | Q | 10.5 |
| Pig | G | K | I | L | K | K | R | R | L | S | L | - | N | Q | 10 |
| Cow | G | K | I | L | K | K | R | R | L | S | L | - | N | Q | 10 |
| Dog | G | K | I | L | K | K | R | R | L | S | L | - | S | Q | 10 |
| Rabbit | G | K | I | L | K | K | R | R | L | S | L | - | N | Q | 9.5 |
| Common vampire bat | G | K | V | L | K | K | R | R | L | S | F | - | N | Q | 10.5 |
| Western European hedgehog | G | K | I | L | K | K | R | R | F | S | L | - | N | Q | 9 |
| African savanna elephant | G | K | I | L | K | K | R | R | L | S | L | - | N | Q | 9.5 |
| Southern two-toed sloth | G | E | V | L | K | K | R | R | L | S | L | - | S | Q | 7 |
| Malayan pangolin | G | K | V | L | K | K | R | R | L | S | L | - | N | Q | 10 |
| Minke whale | G | K | I | L | K | K | R | R | L | S | L | - | N | Q | 10 |
| Nine-banded armadillo | G | K | V | L | K | R | R | R | L | S | L | - | N | L | 11.5 |
| Natal long-fingered bat | G | K | I | L | K | R | R | R | L | T | F | - | N | Q | 10.5 |
| Grey short-tailed opossum | R | K | V | K | K | K | R | R | - | P | F | K | N | H | 8 |
| *NLS mutants* | | | | | | | | | | | | | | | |
| Common bottlenose dolphin | G | K | I | L | K | N | R | W | L | S | L | - | N | Q | <2 |
| Killer whale | G | K | I | L | K | N | R | W | L | S | L | - | N | Q | <2 |
| Naked mole-rat | G | T | V | L | K | K | R | W | L | S | L | - | N | Q | <2 |
| Ord's kangaroo rat | G | K | I | L | K | K | R | L | L | S | L | - | N | Q | <2 |
| Banner-tailed kangaroo rat | G | K | I | L | K | K | R | L | L | S | L | - | N | Q | <2 |
| Southern grasshopper mouse | G | K | I | L | K | K | R | L | L | G | F | - | G | T | <2 |
| Big brown bat | G | K | I | L | K | K | R | W | L | S | F | - | N | Q | <2 |
| American beaver | G | K | I | L | K | K | R | W | L | S | L | - | N | Q | <2 |
| Chinese pangolin | G | K | V | L | K | K | R | W | L | S | L | - | N | Q | <2 |
| Tasmanian devil | K | K | K | E | K | K | K | Q | I | S | E | - | S | H | 3 |
| Koala | Q | E | I | E | K | K | R | Q | L | T | R | - | R | H | <2 |
| Common brushtail | R | K | T | - | E | N | R | R | L | F | V | - | S | H | <2 |
| *Consensus sequence from full alignment* | | | | | | | | | | | | | | | |
| | G | K | I | L | K | K | R | R | L | S | L | - | N | Q | |

KKRR motif amino acid:
- match to human
- synonymous
- non-synonymous to K/R
- non-synonymous

B

| Species | Nucleotide sequence | | | | | | | | | | | | Motif |
|---|---|---|---|---|---|---|---|---|---|---|---|---|---|
| | K | | | K | | | R | | | R | | | KKRR |
| *Intact NLS* | | | | | | | | | | | | | |
| Human | a | a | g | a | a | g | a | g | a | c | g | g | KKRR |
| Chipmanzee | a | a | g | a | a | g | a | g | a | c | g | g | KKRR |
| House mouse | a | a | g | a | a | g | a | g | a | c | g | g | KKRR |
| Norway rat | a | a | g | a | a | g | a | g | a | c | g | g | KKRR |
| Pig | a | a | g | a | a | g | a | g | a | c | g | g | KKRR |
| Cow | a | a | g | a | a | g | a | g | a | c | g | g | KKRR |
| Dog | a | a | g | a | a | g | a | g | a | c | g | g | KKRR |
| Rabbit | a | a | g | a | a | a | a | g | a | c | g | c | KKRR |
| Common vampire bat | a | a | g | a | a | g | a | g | a | c | g | g | KKRR |
| Western European hedgehog | a | a | g | a | a | g | a | g | a | a | g | g | KKRR |
| African savanna elephant | a | a | g | a | a | a | a | g | a | c | g | a | KKRR |
| Southern two-toed sloth | a | a | g | a | a | g | c | g | g | c | g | a | KKRR |
| Malayan pangolin | a | a | g | a | a | g | a | g | a | c | g | g | KKRR |
| Minke whale | a | a | g | a | a | g | a | g | a | c | g | g | KKRR |
| Nine-banded armadillo | a | a | g | a | g | g | a | g | a | c | g | a | KRRR |
| Natal long-fingered bat | a | a | g | a | g | g | a | g | a | c | g | g | KRRR |
| Grey short-tailed opossum | a | a | g | a | a | g | a | g | g | c | g | t | KKRR |
| *NLS mutants* | | | | | | | | | | | | | |
| Common bottlenose dolphin | a | a | g | a | a | c | a | g | a | t | g | g | KNRW |
| Killer whale | a | a | g | a | a | c | a | g | a | t | g | g | KNRW |
| Naked mole-rat | a | a | g | a | a | g | a | g | a | t | g | g | KKRW |
| Ord's kangaroo rat | a | a | g | a | a | g | a | g | a | t | t | g | KKRL |
| Banner-tailed kangaroo rat | a | a | g | a | a | g | a | g | a | t | t | g | KKRL |
| Southern grasshopper mouse | a | a | g | a | a | g | a | g | a | c | t | g | KKRL |
| Big brown bat | a | a | g | a | a | g | a | g | a | t | g | g | KKRW |
| American beaver | a | a | g | a | a | g | a | g | a | t | g | g | KKRW |
| Chinese pangolin | a | a | g | a | a | g | a | g | a | t | g | g | KKRW |
| Tasmanian devil | a | a | g | a | a | g | a | a | a | c | a | a | KKKQ |
| Koala | a | a | g | a | a | g | a | g | a | c | a | a | KKRQ |
| Common brushtail | g | a | a | a | a | t | a | g | a | c | g | a | ENRR |

KKRR motif nucleotide:
- match to human
- synonymous
- non-synonymous to K/R
- non-synonymous

**Supplementary Figure 3. Several mammalian species have acquired mutations in the pro-IL-1α NLS KKRR motif, resulting in predicted loss of nuclear localisation.** (**A**) Pro-IL-1α NLS amino acid sequences of several mammalian species, including those that have lost predicted NLS function. Consensus sequence was taken from the alignment of IL-1α from all isoforms of all 158 species. NLS scores were estimated using NLSmapper. Sequences correspond to $G^{78}$-$Q^{90}$ in human sequence. (**B**) KKRR motif nucleotide sequences for each of the species. Synonymous (green), non-synonymous (red), and non-synonymous to K/R (orange) mutations are highlighted.

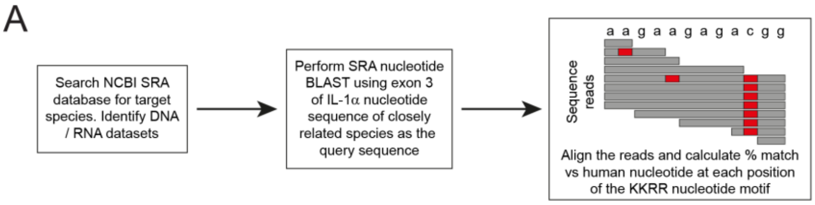


B

| Species | Nucleotide sequence | | | | | | | | | | | | Motif | NLS score |
|---|---|---|---|---|---|---|---|---|---|---|---|---|---|---|
| | K | | | K | | | R | | | R | | | KKRR | |
| American beaver | a | a | g | a | a | g | a | g | a | t | g | g | KKRW | <2 |
| Eurasian beaver | a | a | g | a | a | g | a | g | a | t | g | g | KKRW | <2 |
| Desmarest's spiny pocket mouse | a | a | a | a | a | g | a | g | a | t | g | g | KKRW | <2 |
| Pacific pocket mouse | a | a | g | a | a | g | a | g | a | t | g | g | KKRW | <2 |
| Bailey's pocket mouse | a | a | g | a | a | g | a | g | a | t | g | g | KKRW | <2 |
| Rock pocket mouse | a | a | g | a | a | g | a | g | a | t | g | g | KKRW | <2 |
| Stephen's kangaroo rat | a | a | g | a | a | g | a | g | a | t | t | g | KKRL | <2 |
| Merriam's kangaroo rat | a | a | g | a | a | g | a | g | a | t | t | g | KKRL | <2 |
| Ord's kangaroo rat | a | a | g | a | a | g | a | g | a | t | t | g | KKRL | <2 |
| Banner-tailed kangaroo rat | a | a | g | a | a | g | a | g | a | t | t | g | KKRL | <2 |
| Botta's pocket gopher | a | a | g | a | a | a | a | g | a | t | g | g | KKRW | <2 |

match to human    synonymous    non-synonymous

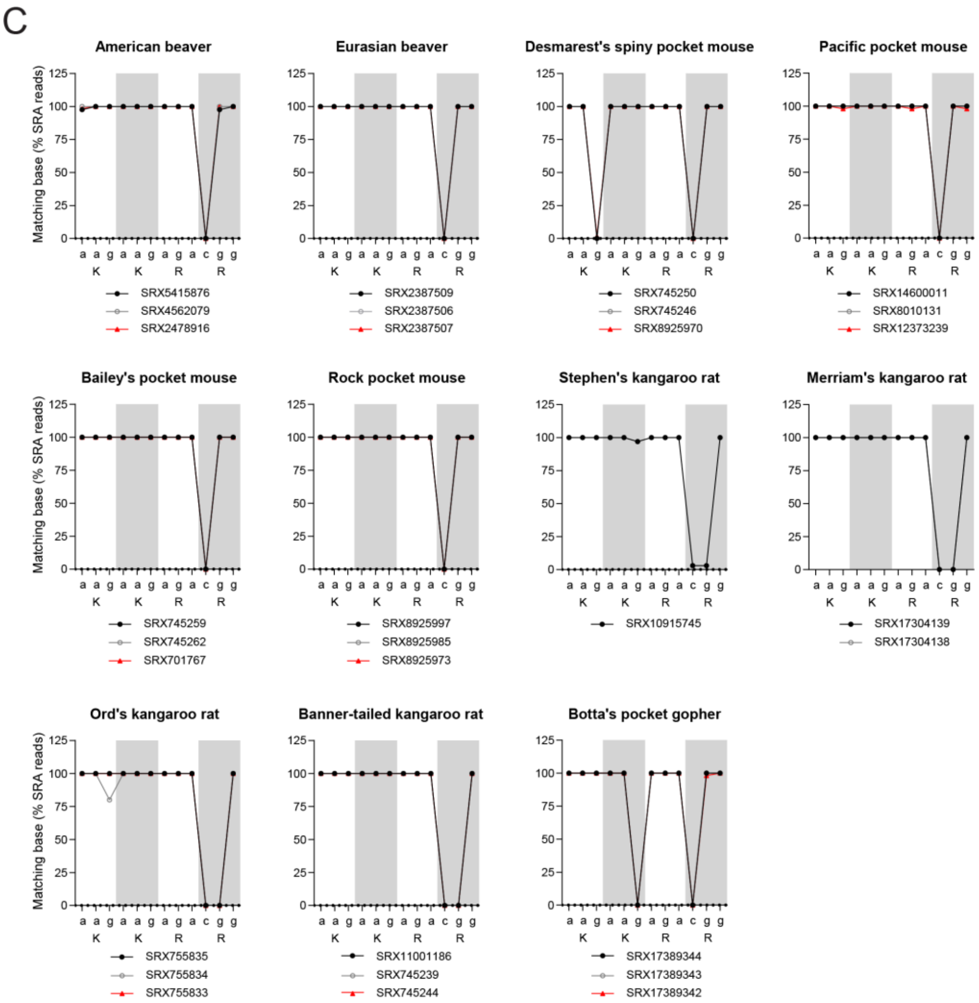


**Supplementary Figure 4. Assessment of NLS mutations in the rodent suborder castorimorpha.** (**A**) Pipeline for retrieving and analysing sequence reads from SRA database to assess KKRR motif nucleotide similarity to human sequence. (**B**) Nucleotide sequences of the KKRR motif of the castorimorpha species, only including species for which an IL-1α sequence could be retrieved. (**C**) SRA reads from the SRA database, which were retrieved by using BLAST with the exon three nucleotide sequence of the American beaver or Ord's kangaroo rat used as the query sequence. Data are presented as % SRA reads that match the human KKRR nucleotide sequence ‘aagaagagacgg’. Where possible, reads were retrieved from at least three individuals, from three separate studies. Synonymous (green), non-synonymous (red), and non-synonymous to K/R (orange) mutations are highlighted.

A

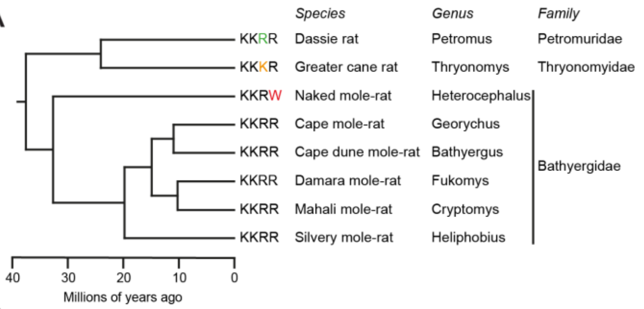


B

| Species | Nucleotide sequence | | | | | | | | | | | | Motif | NLS score |
|---|---|---|---|---|---|---|---|---|---|---|---|---|---|---|
| | K | | | K | | | R | | | R | | | KKRR | |
| Dassie rat | a | a | g | a | a | g | a | g | g | c | g | g | KKRR | 9.5 |
| Greater cane rat | a | a | g | a | a | g | a | a | g | c | g | g | KKKR | 3.5 |
| Naked mole-rat | a | a | g | a | a | g | a | g | a | t | g | g | KKRW | <2 |
| Cape mole-rat | a | a | g | a | a | g | a | g | a | c | g | g | KKRR | 10 |
| Cape dune mole-rat | a | a | g | a | a | g | a | g | a | c | g | g | KKRR | 10 |
| Damaraland mole-rat | a | a | g | a | a | g | a | g | a | c | g | g | KKRR | 10 |
| Mahali mole-rat | a | a | g | a | a | g | a | g | a | c | g | g | KKRR | 10 |
| Silvery mole-rat | a | a | g | a | a | g | a | g | a | c | g | g | KKRR | 9.5 |

match to human synonymous non-synonymous to K/R non-synonymous

C

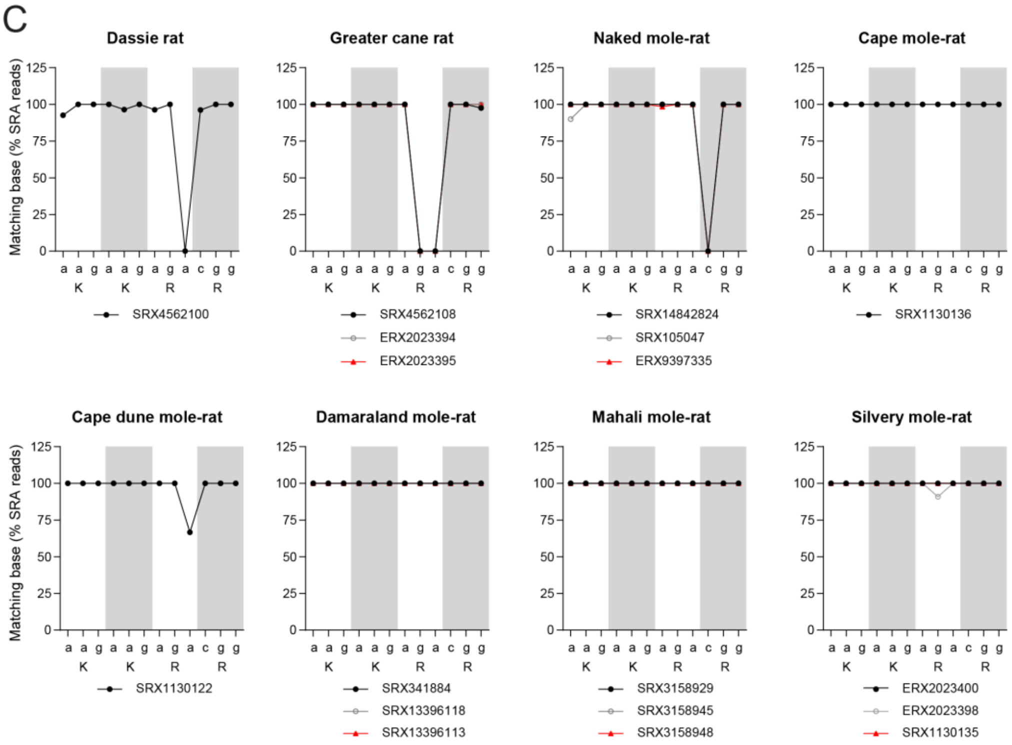


**Supplementary Figure 5. Assessment of NLS mutations in the naked mole-rat superfamily.** (**A**) Model evolutionary tree of the naked mole-rat superfamily, only including species for which an IL-1α sequence could be retrieved. (**B**) Nucleotide sequences of the KKRR motif of these mole-rat species. (**C**) SRA reads from the SRA database, which were retrieved by using BLAST with the exon three nucleotide sequence of the naked mole-rat used as the query sequence. Data are presented as % SRA reads that match the human KKRR nucleotide sequence 'aagaagagacgg'. Where possible, reads were retrieved from at least three individuals, from three separate studies. Synonymous (green), non-synonymous (red), and non-synonymous to K/R (orange) mutations are highlighted. Divergence times were retrieved using TimeTree (15).

A

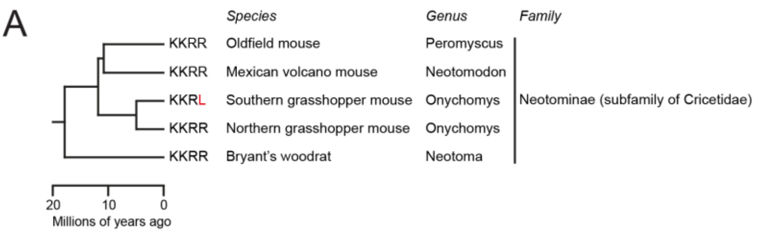


B

| Species | Nucleotide sequence | | | | | | | | | | | | Motif | NLS score |
|---|---|---|---|---|---|---|---|---|---|---|---|---|---|---|
| | K | | | K | | | R | | | R | | | KKRR | |
| Oldfield mouse | a | a | g | a | a | g | a | g | a | c | g | g | KKRR | 10.5 |
| Mexican volcano mouse | a | a | g | a | a | g | a | g | a | c | g | g | KKRR | 10.5 |
| Southern grasshopper mouse | a | a | g | a | a | g | a | g | a | c | t | g | KKRL | <2 |
| Northern grasshopper mouse | a | a | g | a | a | g | a | g | a | c | g | g | KKRR | 9 |
| Bryant's woodrat | a | a | g | a | a | g | a | g | a | c | g | g | KKRR | 9.5 |

match to human    non-synonymous

C

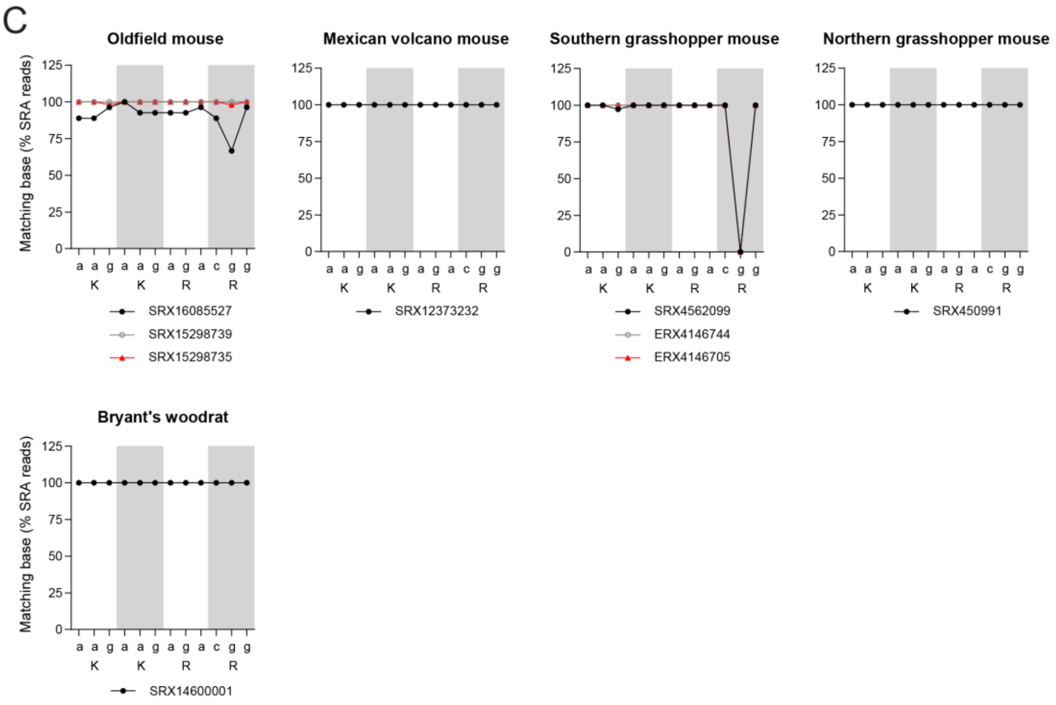


**Supplementary Figure 6. Assessment of NLS mutations in the southern grasshopper mouse family.** (**A**) Model evolutionary tree of the southern grasshopper mouse family, only including species for which an IL-1α sequence could be retrieved. (**B**) Nucleotide sequences of the KKRR motif of these species. (**C**) SRA reads from the SRA database, which were retrieved by using BLAST with the exon three nucleotide sequence of the southern grasshopper mouse used as the query sequence. Data are presented as % SRA reads that match the human KKRR nucleotide sequence ‘aagaagagacgg’. Where possible, reads were retrieved from at least three individuals, from three separate studies. Synonymous (green) and non-synonymous (red) mutations are highlighted. Divergence times were retrieved using TimeTree (15).

A

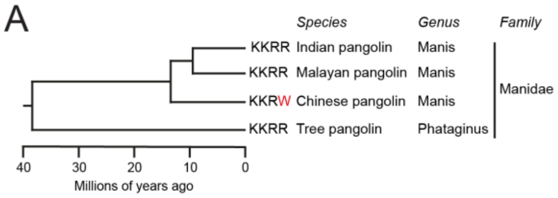


B

| Species | Nucleotide sequence | | | | | | | | | | | | Motif | NLS score |
|---|---|---|---|---|---|---|---|---|---|---|---|---|---|---|
| | K | | | K | | | R | | | R | | | KKRR | |
| Indian pangolin | a | a | g | a | a | g | a | g | a | c | g | g | KKRR | 10 |
| Malayan pangolin | a | a | g | a | a | g | a | g | a | c | g | g | KKRR | 10 |
| Chinese pangolin | a | a | g | a | a | g | a | g | a | t | g | g | KKRW | <2 |
| Tree pangolin | a | a | g | a | a | g | a | g | a | c/a | g | g | KKRR | 10 |

match to human  synonymous  non-synonymous

C

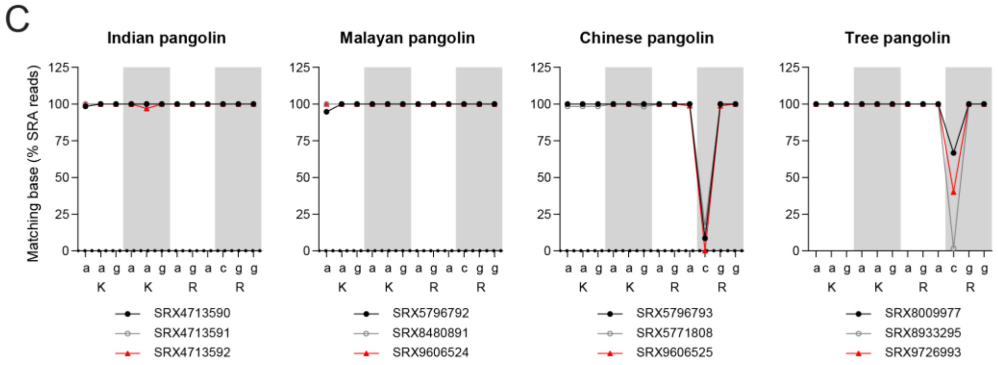


**Supplementary Figure 7. Assessment of NLS mutations in the pangolin family.** (**A**) Model evolutionary tree of the pangolin family. (**B**) Nucleotide sequences of the KKRR motif of these species for which an IL-1α sequence could be retrieved. (**C**) SRA reads from the SRA database, which were retrieved by using BLAST with the exon three nucleotide sequence of the Chinese pangolin used as the query sequence. Data are presented as % SRA reads that match the human KKRR nucleotide sequence 'aagaagagacgg'. Where possible, reads were retrieved from at least three individuals, from three separate studies. Synonymous (green) and non-synonymous (red) mutations are highlighted. Divergence times were retrieved using TimeTree (15).

A

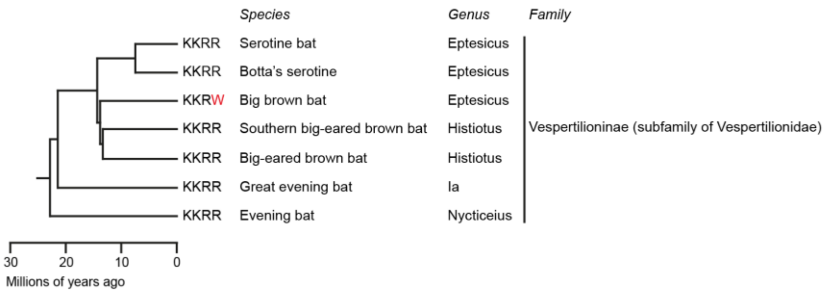


B

| Species | Nucleotide sequence | | | | | | | | | | | | Motif | NLS score |
|---|---|---|---|---|---|---|---|---|---|---|---|---|---|---|
| | K | | | K | | | R | | | R | | | KKRR | |
| Serotine bat | a | a | g | a | a | g | a | g | a | c | g | g/a | KKRR | 10.5 |
| Botta's serotine | a | a | g | a | a | g | a | g | a | c | g | g | KKRR | 14.5 |
| Big brown bat | a | a | g | a | a | g | a | g | a | c/t | g | g | KKRW | <2 |
| Southern big-eared brown bat | a | a | g | a | a | g | a | g | a | c | g | g | KKRR | 11 |
| Big-eared brown bat | a | a | g | a | a | g | a | g | a | c | g | g | KKRR | 11 |
| Great evening bat | a | a | g | a | a | g | a | g | a | c | g | g | KKRR | 10.5 |
| Evening bat | a | a | g | a | a | g | a | g | a | c | g | g | KKRR | 10.5 |

match to human  synonymous  non-synonymous

C

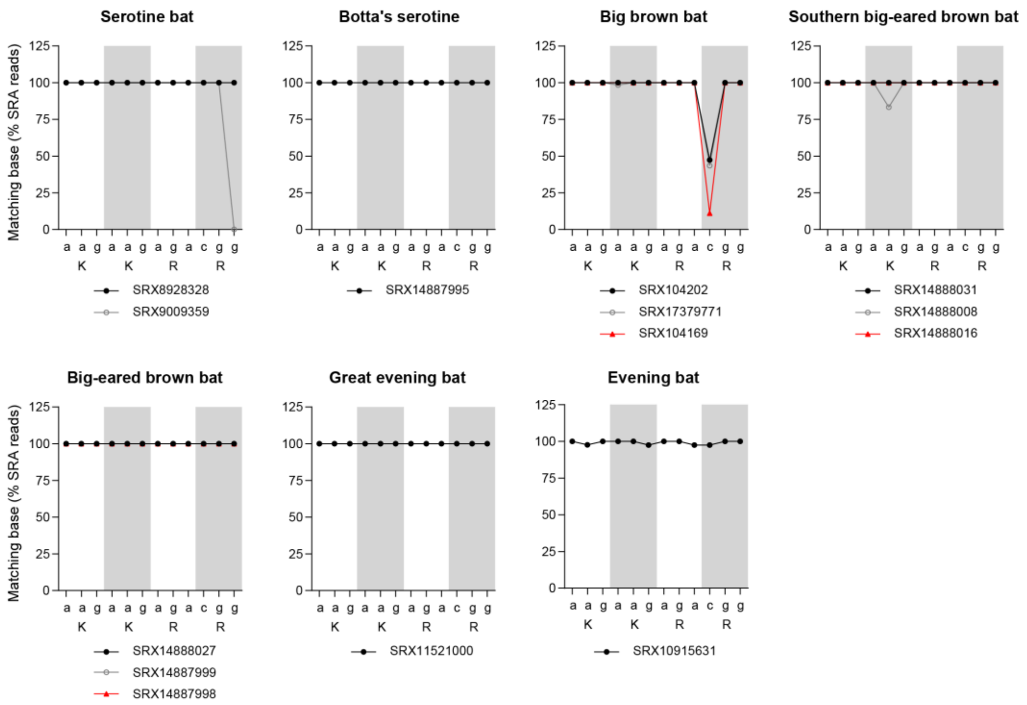


**Supplementary Figure 8. Assessment of NLS mutations in the big brown bat family.** (**A**) Model evolutionary tree of the big brown bat family, only including species for which an IL-1α sequence could be retrieved. (**B**) Nucleotide sequences of the KKRR motif of these species. (**C**) SRA reads from the SRA database, which were retrieved by using BLAST with the exon three nucleotide sequence of the big brown bat used as the query sequence. Data are presented as % SRA reads that match the human KKRR nucleotide sequence 'aagaagagacgg'. Where possible, reads were retrieved from at least three individuals, from three separate studies. Synonymous (green) and non-synonymous (red) mutations are highlighted. Divergence times were retrieved using TimeTree (15).

| Species | Nucleotide sequence | | | | | | | | | | | | Motif | NLS score |
|---|---|---|---|---|---|---|---|---|---|---|---|---|---|---|
| | K | | | K | | | R | | | R | | | KKRR | |
| Grey short-tailed opossum | a | a | g | a | a | g | a | g | g | c | g | t | KKRR | 8 |
| Virginia opossum | a | a | g | a | a | g | a | g | g | t | g | t | KKRC | 6.5 |
| Monito del Monte | a | a | a | g | a | g | a | g | a | a | g | a | KERR | <2 |
| Koala | a | a | g | a | a | g | a | g | a | c | a | a | KKRQ | <2 |
| Common wombat | a | a | g | a | a | g | a | g | a | c | a | a | KKRQ | <2 |
| Coppery ringtail possum | a | a | g | g | a | g | a | g | a | a | g | a | KERR | <2 |
| Western ringtail possum | a | a | g | g | a | g | a | g | a | a | g | a | KERR | <2 |
| Swamp wallaby | a | a | g | a | a | g | a | g | a | c | a | a | KKRQ | <2 |
| Western brush wallaby | a | a | g | a | a | g | a | g | a | c | a | a | KKRQ | <2 |
| Red kangaroo | a | a | g | a | a | g | a | g | a | c | a | a | KKRQ | <2 |
| Western grey kangaroo | a | a | g | a | a | g | a | g | a | c | a | a | KKRQ | <2 |
| Rufous hare-wallaby | g | t | a | a | a | g | a | g | a | c | a | a | VKRQ | <2 |
| Quokka | a | a | g | a | a | g | a | g | a | c | a | a | KKRQ | <2 |
| Yellow-footed rock-wallaby | a | a | g | a | a | g | a | g | a | c | a | a | KKRQ | <2 |
| Matschie's tree-kangaroo | a | a | g | a | a | g | a | g | a | c | a | a | KKRQ | <2 |
| Woylie | a | a | g | a | a | a | a | g | a | c | a | a | KKRQ | <2 |
| Gilbert's potoroo | a | a | g | a | a | g | a | g | a | c | a | a | KKRQ | <2 |
| Common brushtail | g | a | a | a | a | t | a | g | a | c | g | a | ENRR | <2 |
| Ground cuscus | g | a | a | a | a | t | a | g | a | c | g | a | ENRR | <2 |
| Southern marsupial mole | a | a | g | a | a | g | a | g | c | c | a | a | KKSQ | <2 |
| Eastern quoll | a | a | g | a | a | g | a | a | a | c | a | a | KKKQ | <2 |
| Tasmanian devil | a | a | g | a | a | g | a | a | a | c | a | a | KKKQ | 3 |
| Brush-tailed phascogale | a | a | g | a | a | g | a | a | a | c | a | a | KKKQ | <2 |
| Brown antechinus | a | a | g | a | a | g | a | a | a | g | a | a | KKKE | <2 |
| Fat-tailed dunnart | a | a | g | a | a | g | a | a | a | c | a | a | KKKQ | <2 |
| Tasmanian tiger | a | a | g | a | a | g | a | g | a | c | g | a | KKRR | 12 |

match to human

synonymous

non-synonymous to K/R

non-synonymous

**Supplementary Figure 9. Assessment of NLS mutations in the marsupial lineage.** Nucleotide sequences of the KKRR motif of these species. Synonymous (green), non-synonymous (red), and non-synonymous to K/R (orange) mutations are highlighted.

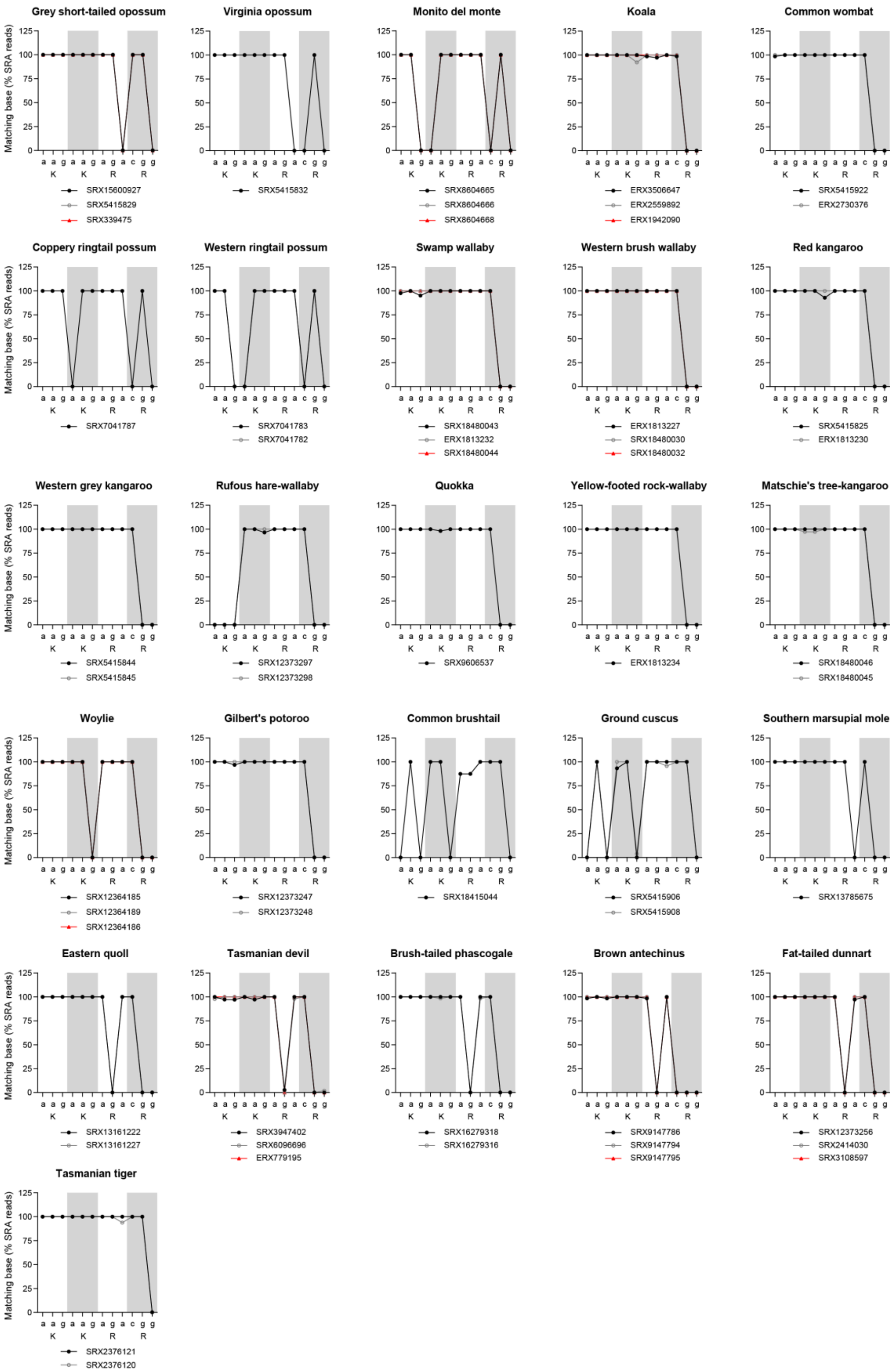


**Supplementary Figure 10. Validation of NLS mutations in the marsupial lineage.** SRA reads from the SRA database, which were retrieved by using BLAST with the exon three nucleotide sequence of the koala, Tasmanian devil or common brushtail used as the query sequence. Data are presented as

% SRA reads that match the human KKRR nucleotide sequence ‘aagaagagacgg’. Where possible, reads were retrieved from at least three individuals, from three separate studies.

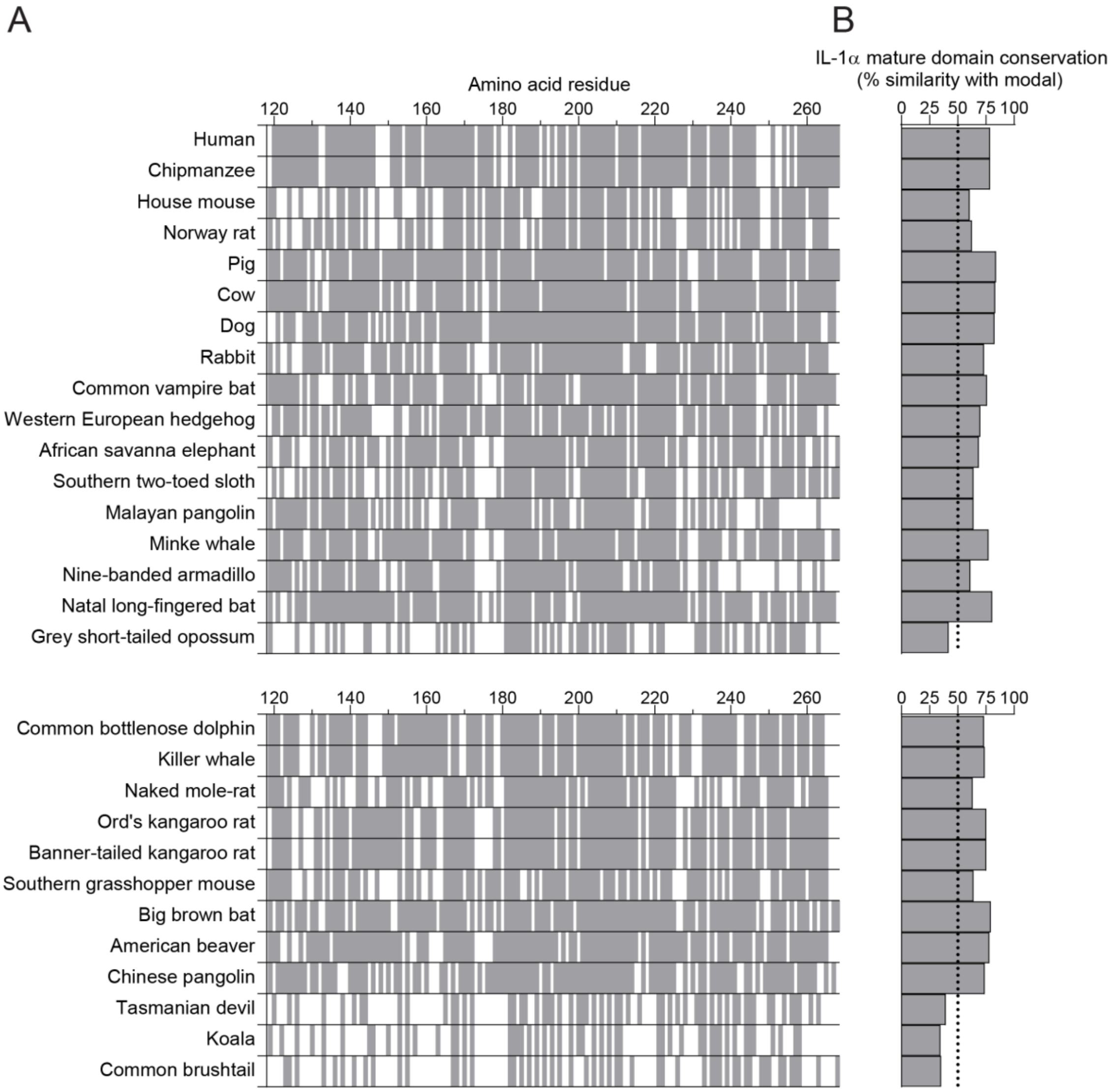


**Supplementary Figure 11. Conservation of the IL-1α mature domain.** (**A**) IL-1α mature domain amino acid conservation in representative intact NLS (top) and mutated NLS (bottom) species. Residues that match the modal amino acid from the full sequence alignment are indicated by a solid grey colour, whereas residues that did not match are indicated by a white gap. Positions where gaps were the modal residue were removed from the alignment. (**B**) Conservation of the whole IL-1α mature domain relative to the modal mature domain sequence.

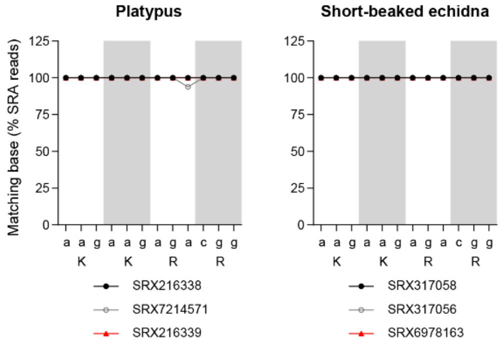


**Supplementary Figure 12. Validation of NLS mutations in the monotreme lineage.** SRA reads from the SRA database, which were retrieved by using BLAST with the exon three nucleotide sequence of the platypus or short-beaked echidna used as the query sequence. Data are presented as % SRA reads that match the human KKRR nucleotide sequence ‘aagaagagacgg’. Where possible, reads were retrieved from at least three individuals, from three separate studies.
